# Cross-talk between intracellular pathogen infection and IFNγ signaling drives enhanced pro-inflammatory responses

**DOI:** 10.64898/2026.05.23.726403

**Authors:** Mullai Valli Ramamoorthy, Bhavyalakshmi Kadavil Baburaj, Devika Jayan, Sandhya Ganesan

**Affiliations:** School of Biology, Indian Institute of Science Education and Research (IISER) Thiruvananthapuram, Maruthamala P.O., Vithura, Thiruvananthapuram, Kerala, India 695551; Friedrich-Alexander-Universitat (FAU) Erlangen-Nurnberg, Erlangen, Germany

## Abstract

Many intracellular pathogens have evolved to evade immune responses and establish a secure niche inside host cells. One such stealth pathogen is the obligate intracellular bacterium *Coxiella burnetii*, the causative agent of Q-fever. *Coxiella* translocates an array of bacterial proteins (‘effectors’) into the host cell through a type IVB secretion system (T4BSS) that mediates suppression of pathogen sensing and innate immunity. Yet, at a systemic level, immunocompetent hosts often restrict pathogens through Th1-mediated and cell-autonomous immunity through the expression of immune-inducible genes. However, the expression and regulation of chemokines, particularly, the CXC-ligands (CXCL9,-10,-11) that are considered biomarkers of Q-fever, is poorly understood. We observed minimal to no *CXCL10* transcript levels during *Coxiella* infection. However, *Coxiella*-infected cells robustly augmented IFNγ-activated expression of CXCL10 in both phagocytic and non-phagocytic cells, and this process was dependent on viability and T4BSS in epithelial cells. This phenomenon extends to other highly pro-inflammatory cytokines and other pathogens including *Salmonella, Mycobacteria* (H37Ra) and *Toxoplasma*. Synergistic increase in *CXCL10* expression in *Coxiella*-infected, IFNγ-activated cells requires ISRE and NF-κB transcriptional elements in the promoter, and the transcription factors STAT1, STAT3 and IRF9. Inhibition of STAT3 by small molecule inhibitors potently decreased the excess promoter activity of *CXCL10*. In addition, treatment of *Coxiella*-infected cells with IFNγ is associated with decreased expression of *SOCS1*, a negative regulator of the IFNγ signaling axis and relatively higher detection of extracellular bacteria. Altogether, these data demonstrate that intracellular pathogens including those conventionally considered to be “immunologically silent”, robustly synergize with IFNγ signaling, with STAT3 activation emerging to be a nodal point for promoting both persistent infection as well as synergism in the expression of immune genes.

**Author summary:** Acute host immune response is often associated with production of soluble messenger molecules called cytokines/chemokines which direct the migration, recruitment and activation of leukocytes and serve as biomarkers in infectious and inflammatory diseases. The regulation of expression of these molecules and their influence on the infection process is not well-understood. In particular, interferon-gamma (IFNγ), a potent pro-inflammatory cytokine produced by activated T and NK cells, activates signaling pathways involved in host defense and inflammation in macrophages and other cell types. We observed that infection with many intracellular bacterial/parasitic pathogens that employ sophisticated immune evasion strategies, synergize with IFNγ signaling and significantly amplify the levels of pro-inflammatory mediators implicating the origin of adverse immune pathologies. We investigated the mechanistic basis of this seemingly counter-intuitive phenomenon, underlying host and bacterial factors involved in distinct cell types, and identified the small molecule-targetable-transcription factor STAT3 as a host determinant in promoting excess cytokine synthesis.

## Introduction

Activation of innate immunity and secretion of cytokines and chemokines that promote chemotaxis, proliferation, activation of immune cells is critical for mounting an effective primary defense against microbial infections. However, many “stealth” pathogens have evolved nuanced mechanisms to replicate inside hostile subcellular compartments and evade immune surveillance and hence, pose a challenge for effective detection and clearance. These pathogens deploy a wide arsenal of effectors and employ several strategies, including inhibiting apoptosis, subverting autophagy, modulation of PRR signaling pathways and host cell membrane traffic etc. to persist and replicate inside host cells [1–3]. Consequently, such mammalian-adapted pathogens have served as excellent models to unveil layered mechanisms of pattern, stress and danger recognition by the host immune system.

The Q fever-causing, gram-negative obligate intracellular bacterium *Coxiella burnetii* is a particularly interesting model of host-pathogen interaction due to its high infectivity, its zoonotic and epidemic potential, Category B select agent status, and obligate intracellular and lysosome-adapted lifestyle [4–6]. *C. burnetii* is largely considered an immunologically silent pathogen owing to the (i) lack of an immunostimulatory hexaacylated LPS and flagellin, (ii) shielding of surface lipoproteins by O-antigen, and (iii) inhibition of infection-induced apoptosis [7–12]. Further, an array of *Coxiella* effectors secreted through its type IVB secretion system (T4BSS) facilitate evasion or inhibition of various innate immune signaling pathways (example: RIG-I, inflammasome) and activation of key transcription factors (example: NF-κB, IRF3) interfering with the upregulation of cytokines and genes with antimicrobial functions [13–18]. However, the adaptive immune response, particularly the T cell immunity and its effector functions, play an essential role in the clearance of intracellular *C. burnetii* infection as evidenced by investigating the effect of deficiencies in MHC-I and II, T cells and IFNγ receptor (IFNγR) [19–21]. The pro-inflammatory cytokine, IFNγ, produced by activated T and NK cells, controls many viral, bacterial, fungal and parasitic pathogens through robust upregulation of Interferon Stimulated Genes (ISGs) [22]. IFNγ signals through the JAK-STAT pathway by activating JAK1 and JAK2 kinases, which phosphorylate STAT1 and promote its dimerization and nuclear translocation. Inside the nucleus, STAT1 cooperates with transcription factors to drive the expression of ISGs by binding to gamma-activated sequence (GAS) elements in their promoters. Our work demonstrated the potency of IFNγ pathway in restricting intracellular *C. burnetii* replication in a cell-autonomous manner and the requirement of the ISGs, Indoleamine dioxygenase 1 (*IDO1*) and Syntaxin 11 (*STX11*) [23,24]. However, the full spectrum of IFNγ-induced genes and their precise roles in pathogen restriction, immune regulation, and disease pathology remain incompletely understood. Among the most highly induced interferon-stimulated genes (ISGs) are the CXCL-family chemokines CXCL9, CXCL10, and CXCL11, which are increasingly recognized for their immunomodulatory functions [25,26]. While there has been a growing understanding of the important roles of type I interferons and ISGs in bacterial infections, the role and regulation of chemokines, particularly in the disease pathogenesis of *Coxiella* and many other intracellular pathogens is not well-defined. Clinical evidence highlights a consistent association between CXCL-chemokines and Q fever pathophysiology. Transcriptomic analyses of peripheral blood mononuclear cells (PBMCs) stimulated with *C. burnetii,* together with serum protein analyses, show marked upregulation of CXCL9, CXCL10, CXCL11 in chronic Q fever patients compared to healthy controls [27]. Similarly, stimulation of whole blood from patients with Q fever fatigue syndrome (QFS) exhibit significantly elevated IFNγ and CXCL10 responses relative to recovered individuals [28]. Enhanced IFNγ production is also observed in PBMCs from chronic Q fever patients following restimulation with *C. burnetii* strains, with CXCL9, CXCL10, and CXCL11 among the most highly induced genes, but not after unrelated stimuli such as *E. coli* LPS [29]. Consistent with these findings, serum CXCL10 levels positively correlate with IFNγ in chronic Q fever [30]. Elevated IFNγ, CXCL10, and IL-2 responses have additionally been reported in individuals previously exposed to *C. burnetii*, through natural infection or vaccination [31,32]. Collectively, these findings suggest that a high IFNγ/IL-2 ratio, together with increased CXCL chemokine expression, may serve as potential biomarkers for chronic Q fever and reflect sustained immune activation associated with disease progression [27,32]. Altogether, these data show an enhanced capacity of PBMCs from chronic Q fever- infected or *Coxiella burnetii*- exposed individuals to produce CXCL chemokines upon re-stimulation suggesting a state of altered immune programming in these cells. Further, the positive correlation between CXCL-chemokines and IFNγ levels indicates that elevated IFNγ alone is insufficient to restrict infection. Importantly, this observation indicates that *C. burnetii* may not actively suppress CXCL chemokine production, in contrast to the immune evasion strategies commonly employed by other intracellular pathogens that target innate signaling pathways. These findings prompted us to investigate the molecular mechanisms regulating CXCL chemokine synthesis during infection and in response to IFNγ stimulation.

This study focuses on CXCL10 (IP-10), a 10 kDa pro-inflammatory chemokine that signals through the CXCR3 receptor to recruit immune cells to infection sites [33,34]. Although primarily associated with IFNγ induction, CXCL10 expression is also driven by Type I IFNs and other cytokines as well as PRR signaling. Beyond its role in chemotaxis, CXCL10 also regulates cell growth and proliferation, as well as apoptosis and angiogenesis [34]. In this work, we utilize *Coxiella burnetii* as a stealth pathogen model to define the molecular drivers of CXCL10 production and its subsequent effect on intracellular replication. We further characterize the bacterial and host factors governing this response and assess its broader relevance across various pathogens, cytokines and cell types.

## Results

### *C. burnetii* augments IFNγ-induced expression of *CXCL10*

We first set out to determine whether *C. burnetii* induces the expression of *CXCL10* in non-phagocytic cells (Figure 1A) and phagocytic cells (Figure 1B). Studies involving human PBMCs typically utilize samples from patients infected with virulent, O-antigen-expressing strains or employ them for re-infection [27,29]. In contrast, the BSL2-adapted NMII phase variant lacks the O-antigen and outer LPS core, thereby exposing surface lipoproteins to host PRRs and allowing dendritic cell maturation [8,9,35,36]. Because NMII is a more potent activator of innate immunity and replicates intracellularly with similar kinetics as NMI, we used this strain to elicit higher baseline immune responses [8,37].

**Figure 1.**
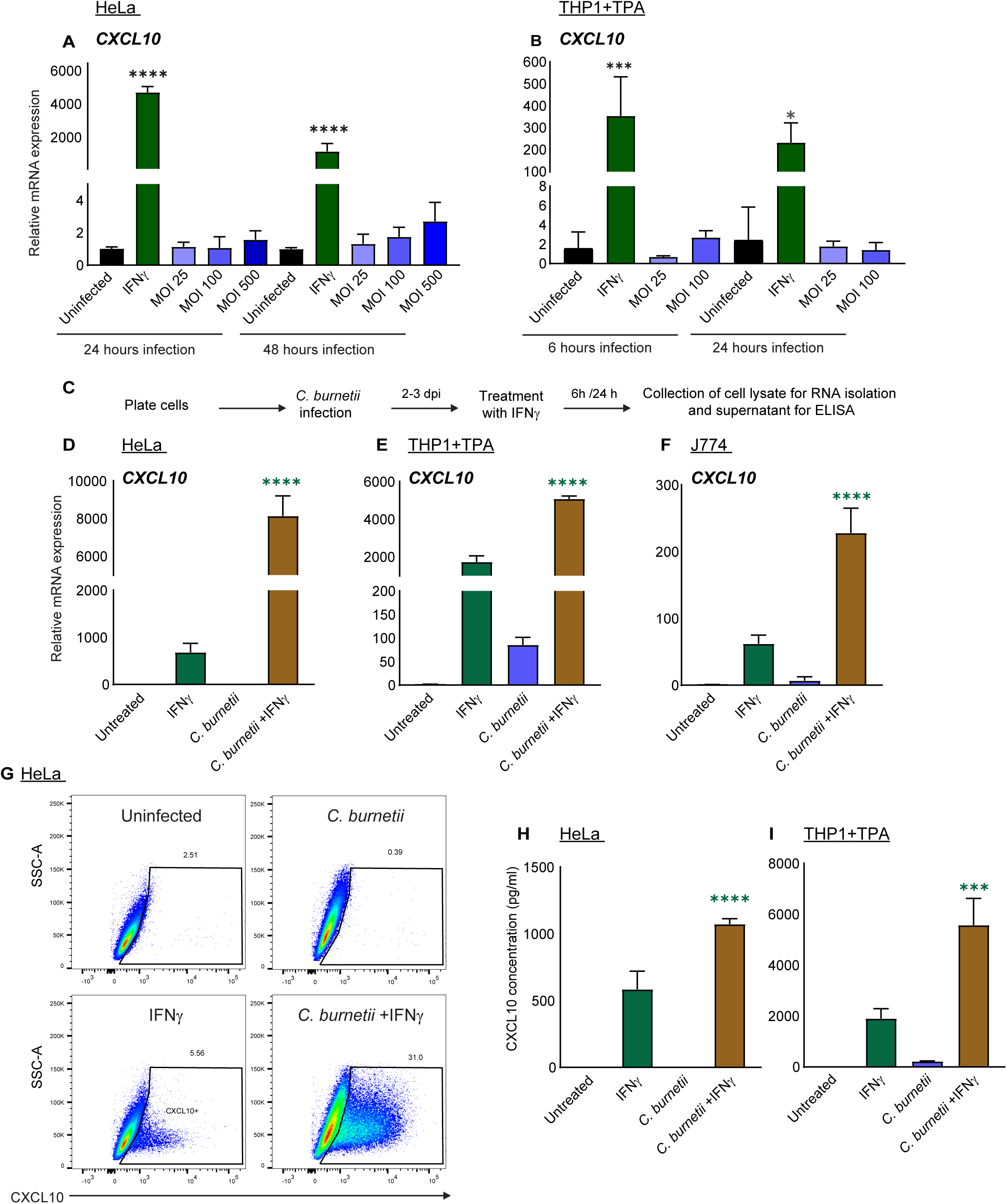
*C. burnetii* augments IFNγ-induced expression of *CXCL10*. Relative mRNA expression of *CXCL10* in HeLa (**A**) and TPA-treated THP1 macrophages (**B**) infected with *C. burnetii* NMII at various MOI (Multiplicity Of Infection) or time points as indicated. Experimental layout for panels **D**, **E**, **F**, **H** and **I** is given in **C**. Relative mRNA expression of *CXCL10* in HeLa (**D**), THP1 macrophages (**E**), and J774 macrophages (**F**) infected with *C. burnetii* followed by supplementation with recombinant human or mouse IFNγ (10 ng/ml for HeLa and 100 ng/ml for THP1 and J774 cells) (hereafter referred to as “synergistic condition”). Protein concentration of CXCL10 in the supernatants of HeLa (**H**) or THP1 macrophages (**I**) in the synergistic conditions. Samples were collected for RNA isolation and ELISA after 24 hours of IFNγ treatment for HeLa and 6 hours for THP1 and J774 cells. Values indicate mean with SD and are the representative of three independent experiments (X=3, n=3). The values are normalized relative to GAPDH expression (**A, B, D-F**). Statistical analysis was performed using one-way ANOVA with Tukey’s post-hoc test. * p<0.05, *** p<0.001, **** p<0.0001 compared to uninfected (black bar), for panels A and B and *** p<0.001, **** p<0.0001 compared to IFNγ-alone treatment (green bar), for panels D, E, F, H and I. In panel **G**, HeLa cells were either uninfected or infected with *C. burnetii* for 3 days followed by treatment with or without IFNγ (10 ng/ml) for 24 hours as depicted in panel C. Sixteen hours post IFNγ treatment, cells were treated with Monensin (0.67 µl per million cells/ml), stained for CXCL10 and measured by flow cytometer to assess intracellular CXCL10 protein levels (**G**). The graph is the representative of three independent experiments (X=2, n=3).

*C. burnetii* induced minimal to no *CXCL10* mRNA levels even at the highest MOI used (Figure 1A and 1B). To verify productive infection, genomic equivalents were measured through qPCR using *Coxiella*-specific *dotA* primers, which showed an increase over time and MOI (Figure S1A and S1B). CXCL10 is a well-established, highly inducible transcriptional target of IFNγ signaling [33,38], hence, activation by IFNγ served as a positive control (Figure 1A and 1B). Importantly, *CXCL10* induction was also less-pronounced in the phagocytic, TPA-differentiated human macrophage line (THP1) infected with *C. burnetii* which are equipped with a broader spectrum of microbial pattern recognition pathways (Figure 1B). It is possible that *Coxiella* is a poor inducer of *CXCL10* due to a failure in generating the PAMPs required to engage its transcriptional pathways. These results also raised the question whether *Coxiella* may encode for mechanisms to actively inhibit *CXCL10* expression. Indeed, low CXCL10 and high transglutaminase 2 (TGM2) levels have been observed in human alveolar macrophages infected with *C. burnetii* suggesting a switch from M1 to M2 state [39]. To address this possibility, we infected HeLa, THP1 human macrophages and J774 mouse macrophages for 2-3 dpi to allow the translocation of effectors and generation of a mature *Coxiella*-containing vacuole (CCV) and subsequently treated them with IFNγ for 24h and 6h for HeLa and macrophages respectively (Figure 1C). Interestingly, *Coxiella* infection significantly augments IFNγ-induced *CXCL10* expression and synergizes with IFNγ signaling in all the cell types examined (Figure 1D-F). The synergistic increase in *CXCL10* transcript levels was also reflected in the levels of the CXCL10 protein synthesized intracellularly as well as that secreted into the supernatants as detected by flow cytometry (Figure 1G) and sandwich-ELISA respectively (Figure 1H and 1I). The percentage of cells positive for CXCL10, as detected by intracellular staining using anti-CXCL10 antibody was higher in *C. burnetii-*infected, IFNγ-treated cells compared to cells treated with IFNγ or *C. burnetii* alone (Figure 1G). We observed that *Coxiella* infection alone resulted in decreased protein production compared to that of the uninfected cells, which is in alignment with a previous study [39], where primary human alveolar macrophages produced less CXCL10 upon *C. burnetii* infection.

Assuming all cells are activated by IFNγ, we tested whether the enhanced expression of CXCL10 originated from infected or by-stander cells (Figure S1C). Flow cytometry analysis using GFP-expressing *Coxiella* and gating for GFP+ population showed that the increased CXCL10 production was primarily associated with infected cells (Figure S1C). Together, these results showed that *C. burnetii* infection does not suppress IFNγ-induced CXCL10, instead it synergizes with the IFNγ-JAK-STAT pathway to augment CXCL10 synthesis and secretion in infected cells. To understand if the increase in the CXCL10 expression depends on the concentration of IFNγ, we treated THP1 macrophages with increasing doses of IFNγ in synergistic conditions (as laid out in Figure 1C). We observed a dose-dependent synergistic increase in the range of 1-100 ng/ml (Figure S1D).

### Synergistic increase in *C. burnetii* and IFNγ-induced *CXCL10* requires bacterial viability and T4BSS activity

To better understand the mechanistic basis of infection +IFNγ induced synergistic increase in *CXCL10* expression (Figure 1D-F), we set out to determine the bacterial and host determinants underlying this phenomenon. When HeLa cells were treated with paraformaldehyde-fixed or heat-killed *C. burnetii* prior to treatment with IFNγ, the synergistic increase in *CXCL10* was significantly abolished (Figure 2A-B). A complete abrogation in synergy demonstrated that bacterial viability is critical. We then questioned whether bacterial protein synthesis was required. Addition of chloramphenicol, that inhibits bacterial protein synthesis by binding to the 50S subunit of prokaryotic ribosomes, to *Coxiella*-infected cells, significantly decreased *CXCL10* expression in infected, IFNγ-treated cells (Figure 2A). Inhibition of bacterial protein synthesis also affects production and secretion of effectors through the T4BSS and thereby, intracellular replication and development of the mature CCV. We tested whether T4BSS activity and productive infection was required for synergistic expression of *CXCL10* by utilizing the *icmL*::Tn *C. burnetii*, that is deficient in the assembly of a functional T4BSS apparatus [40–42] (Figure 2C). The phenotype was observed only with the wild-type *C. burnetii* but not with the *icmL*::Tn mutant, similar to the case with heat-killed/PFA-fixed/chloramphenicol-treated *C. burnetii* indicating the importance of a productive infection and possibly effects of single or multiple effector functions in driving the synergistic expression of *CXCL10* in IFNγ-treated HeLa cells. The phenotype was also evident at the secreted protein level as measured by ELISA (Figure 2D). Contrary to what we observed in HeLa cells, neither bacterial viability nor T4BSS activity was required for *C. burnetii* to increase the expression of *CXCL10* in IFNγ-treated THP1 macrophages both at the protein (Figure S2A) or mRNA (Figure S2B and S2C) levels.

**Figure 2.**
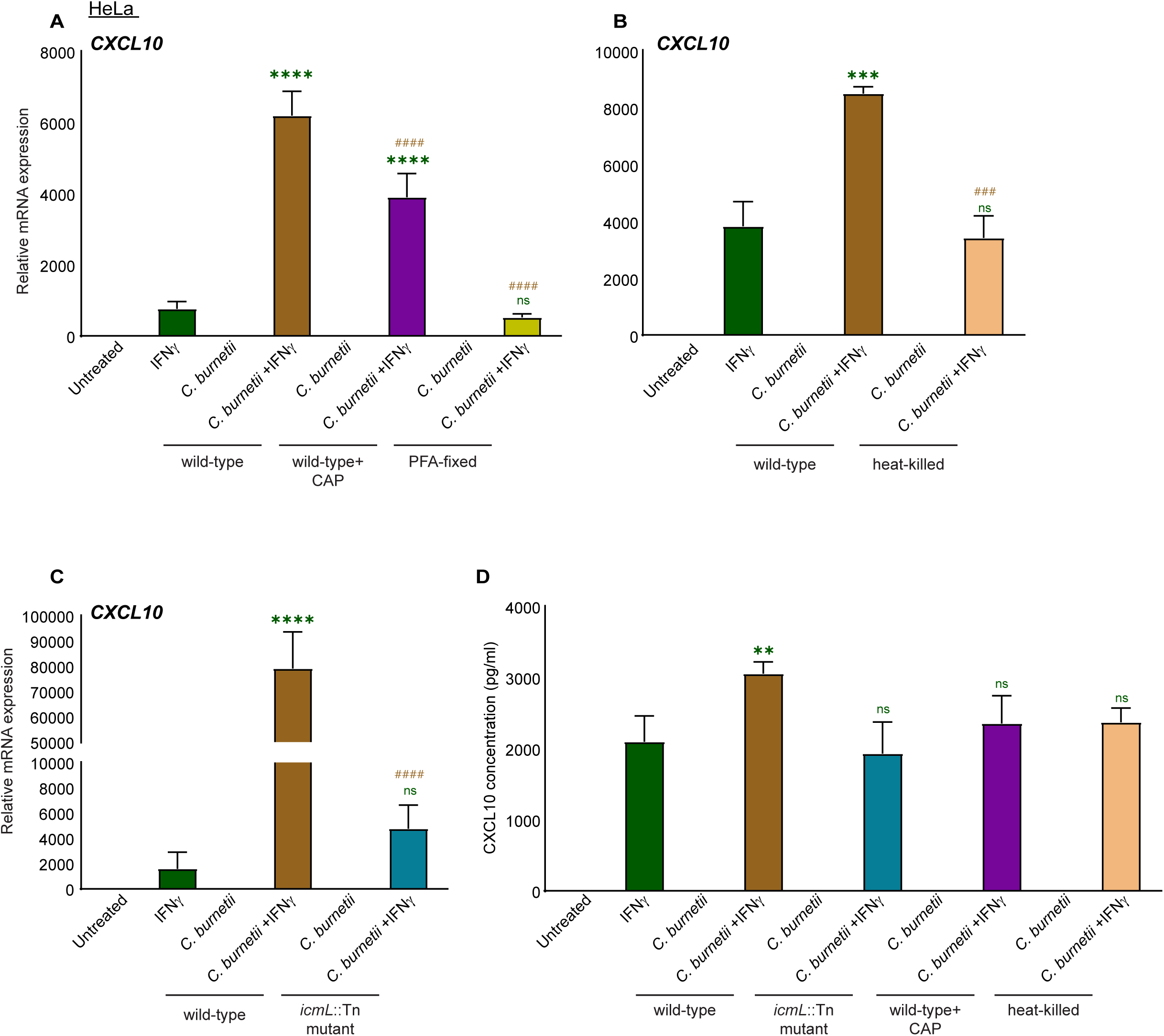
Synergistic increase in *C. burnetii* and IFNγ-induced *CXCL10* requires bacterial viability and T4BSS activity. Relative mRNA expression of *CXCL10* in HeLa cells infected with either wild-type *C. burnetii* (followed by treatment with chloramphenicol (CAP) (10 µg/ml) where indicated) or PFA-fixed (**A**), heat-killed (**B**), or *icmL*::Tn mutant (**C**) followed by treatment with IFNγ, 3 or 4 days post-infection for 24 hours. Concentration of secreted CXCL10 protein as measured by ELISA (**D**). Values indicate mean with SD and are the representative of three independent experiments (X=3, n=3). The values are normalized relative to GAPDH expression (**A-C**). Statistical analysis was performed using one-way ANOVA with Tukey’s post-hoc test. ** p<0.01, *** p<0.001, **** p<0.0001 compared to IFNγ-alone treatment (green bar), ^###^ p<0.001, ^####^ p<0.0001 compared to synergistic conditions with wild-type *C. burnetii* (*C. burnetii* +IFNγ, as represented by the brown bar).

### ISRE and NF-κB transcriptional elements contribute to enhanced IFNγ-induced *CXCL10* in *C. burneti*-infected cells

CXCL10 promoter is transcriptionally activated by multiple transcription factors and signaling pathways [43–48]. In order to understand the mechanistic basis of the synergistic expression of *CXCL10*, we set out to identify the relative contribution of specific transcriptional elements and possibly uncouple elements required for inducing *CXCL10* by IFNγ alone versus that induced by infection +IFNγ. We utilized firefly luciferase-based *CXCL10* reporter constructs where the luciferase expression was driven by either the full length (CXCL10-GL4), truncated (tCXCL10-GL4) or variants of the *CXCL10* promoter with binding sites for specific transcription factors - (NF-κB1, NF-κB2, CCAAT-enhancer-binding protein (CEBPβ1 and -β2) or interferon-stimulated response elements (ISRE), (details given in Table 3) mutated (Figure 3A-E) [46]. The firefly luciferase activity of full-length *CXCL10* promoter was significantly higher in cells infected with *Coxiella* and subsequently activated by IFNγ compared to that by IFNγ alone (Figure 3B), similar to what has been observed with *CXCL10* transcript levels indicating that the enhanced expression originates from differential promoter activity or a level upstream in the pathway. In addition to the full length promoter, cells were transfected with the truncated tCXCL10-GL4 plasmid that lacks the region upstream of −279 of the promoter including the two sites corresponding to binding CEBPβ (Figure 3C and 3D). Synergistic increase in luciferase activity remained unaffected in the case of truncated as well as both CEBPβ mutants (Figure 3C and 3D) indicating that the −1 to −279 proximal, cis-element was sufficient for synergistic expression. However, the synergism was significantly decreased when cells were transfected with constructs that harbor mutations in either the κB1 or κB2 sites and completely abrogated in the construct harboring mutated ISRE site (Figure 3E). Notably, ISRE site was required for luciferase activity not only by infection +IFNγ but also by IFNγ alone indicating that the engagement of this site is critical for IFNγ-induced transcriptional activation of *CXCL10* and a prerequisite for synergistic increase in luciferase activity. Altogether, these data indicate that the NF-κB binding sites are required, particularly for infection-associated synergistic increase in *CXCL10* promoter activity.

**Figure 3.**
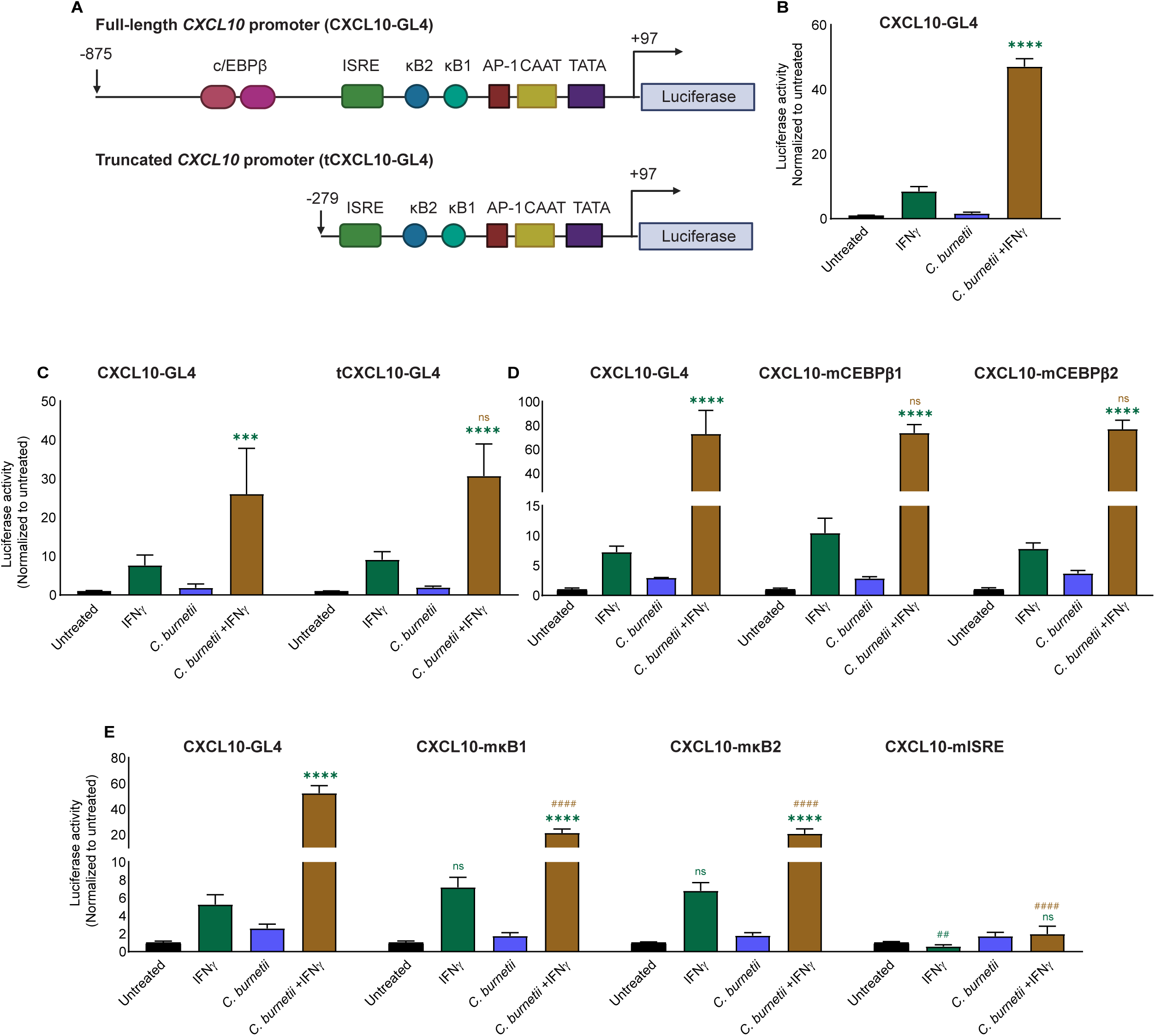
ISRE and NF-κB transcriptional elements contribute to enhanced IFNγ-induced *CXCL10* in *C. burnetii*-infected cells. Schematic representation of the constructs expressing luciferase under the control of full length or variants of human *CXCL10* promoter (**A**). Luciferase activity of *C. burnetii*-infected HeLa cells transfected with full-length (CXCL10-GL4) (**B**), truncated (tCXCL10-GL4) (**C**), or constructs with mutated CEBPβ-binding sites (CXCL10-mCEBPβ1 and CXCL10-mCEBPβ2) (**D**), mutated NF-κB-binding sites (CXCL10-mκB1 and CXCL10-mκB2) and mutated ISRE-binding sites (CXCL10-mISRE) (**E**) at 2 dpi and treated with IFNγ at 3 dpi for 24 hours. Values indicate mean with SD and are representative of three independent experiments (X=4, n=3). Statistical analysis was performed using two-way ANOVA with Tukey’s post-hoc test. *** p<0.001, **** p<0.0001 compared to IFNγ-alone treatment (green bar) of the corresponding CXCL10 promoter construct, ^####^ p<0.0001 compared to synergistic conditions (*C. burnetii* +IFNγ, brown bar) using full-length CXCL10-GL4. In addition, for panel **E**, statistical differences comparing IFNγ-alone treatments corresponding to variant promoter constructs with that of CXCL10-GL4 were indicated by ^##^ p<0.01 or ns.

### STAT1 and IRF9 are required for synergistic increase in *CXCL10* expression in *C. burnetii*-infected cells treated with IFNγ

We hypothesised that the increase in the expression of IFNγ-induced *CXCL10* in the *C. burnetii*-infected cells could, at least in part, be due to 2 possibilities. (i) Infection positively influences IFNγ signaling, (ii) IFNγ-mediated intracellular bacterial killing/ restriction leads to higher PRR signaling likely through generation of more bacterial PAMPs. In addition, our luciferase reporter-based assays show the requirement of NF-κB in the synergistic activation of the promoter (Figure 3E). We tested these possibilities categorically as detailed in the following sections.

To address the first possibility, we set out to identify the transcription factors that potentially contribute to synergistic increase of *CXCL10* including STAT1, STAT2, and selected interferon regulatory factors (IRFs), specifically IRF3, IRF7, and IRF9 which are established as key mediators of interferon-stimulated gene (ISG) induction [49–54]. In particular, the transcription factors STAT1, STAT2, and IRF9 are known to form the interferon-stimulated gene factor 3 (ISGF3) complex that is activated in response to type I interferon signaling, bind to ISRE in the promoters of interferon-stimulated genes (ISGs) to drive their expression [49,50,52]. IFNγ also has been reported to induce the formation of ISGF3 [55,56]. Hence, we transfected the infected cells with the transfection reagent alone (Mock), a non-targeting control siRNA (NTC) or siRNA targeting STAT1, STAT2, IRF3, IRF7 or IRF9 (details given in Table 4) along with the full length CXCL10 promoter construct (CXCL10-GL4) and measured *CXCL10* promoter activity (Figure 4A-C). Synergistic increase in *CXCL10* expression was found to be dependent on IRF9 and STAT1 but does not require STAT2, IRF3 or IRF7 (Figure 4B and 4C). While the possibility of inefficient STAT2 knockdown can not be ruled out, this data appears to point to a non-canonical, STAT2-independent ISGF3 variant that drives IFNγ-induced induction of *CXCL10* [57].

**Figure 4.**
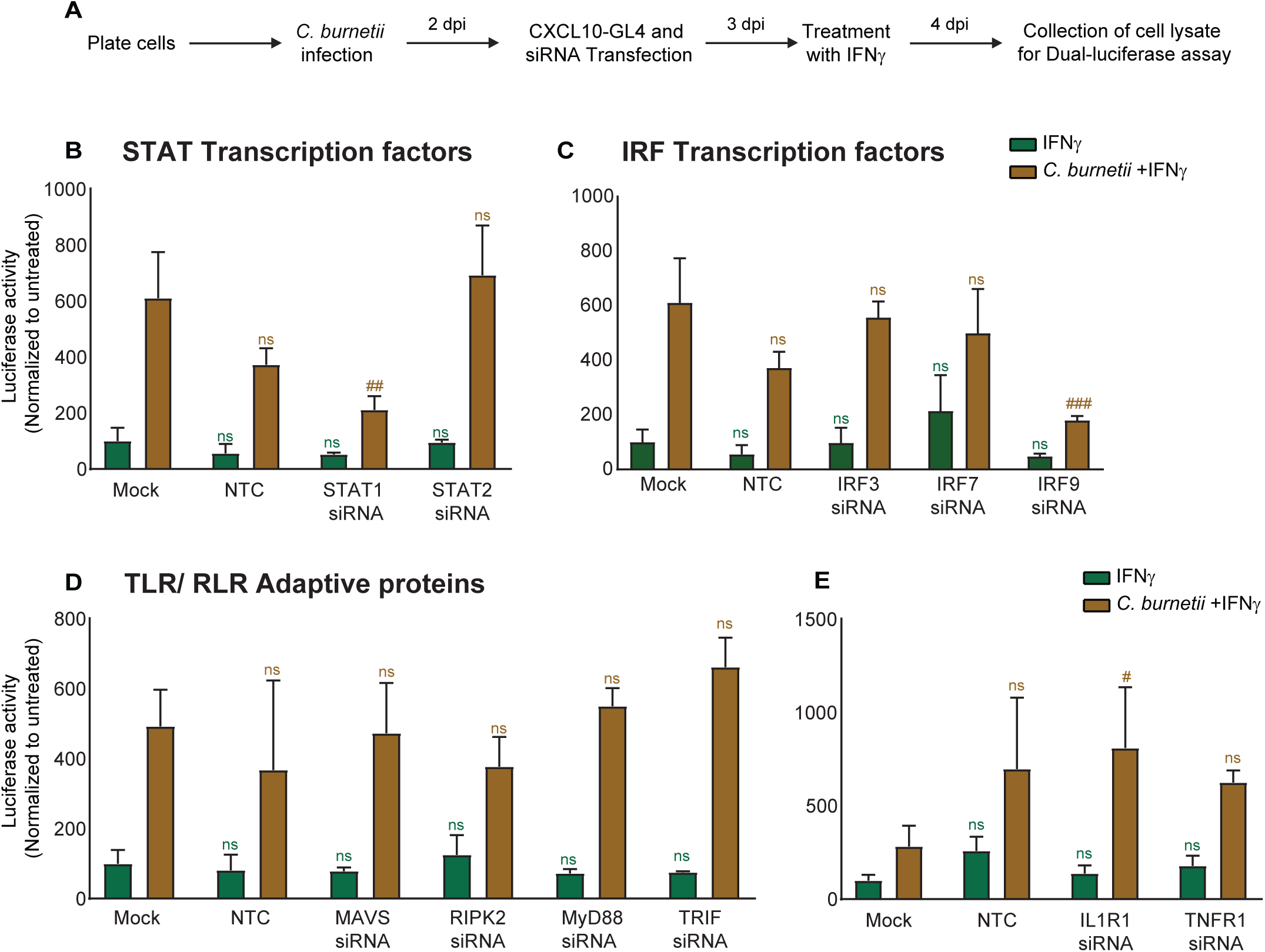
STAT1 and IRF9 are required for synergistic increase in *CXCL10* expression in *C. burnetii*-infected cells treated with IFNγ. Experimental layout is given in **A**. Luciferase activity of *C. burnetii*-infected HeLa cells transfected with full length CXCL10 promoter construct (CXCL10-GL4) along with siRNAs targeting STAT/IRF transcription factors, adaptor proteins of TLR/RLR/NLR signaling pathways or cytokine receptors (25 nM) as indicated (**B-E**) on 2 dpi and treated with IFNγ (10 ng/ml) at 3 dpi for 24 hours. Values indicate mean with SD and are representative of three independent experiments (X=3, n=3). Statistical analysis was performed using two-way ANOVA with Tukey’s post-hoc test. ^#^ p<0.05, ^##^ p<0.01, ^###^ p<0.001 compared to mock-transfected, infected cells treated with IFNγ (*C. burnetii* +IFNγ).

The pattern recognition receptors TLR2, TLR4, NOD2, and TLR1/TLR2 heterodimer and RIG-I have been implicated in the innate immune sensing of *Coxiella* and cytokine production [7,21,58–61]. In addition, *Coxiella* has been demonstrated to inhibit RIG-I signaling pathway by T4BSS-secreted effectors, to reduce type I IFN response [16]. To address the second hypothesis that activation of infected cells with IFNγ leads to the exposure of more PAMPs which can in turn activate various PRR signaling pathways, we tested the requirement of distinct innate immune sensing pathways, by individually silencing the key adaptor proteins MAVS (the mitochondrial antiviral-signaling adaptor downstream of the RIG-I like Receptors RIG-I/MDA5), RIPK2 (a kinase adaptor shared by the NOD-like Receptors NOD1/NOD2), MyD88 (the canonical Toll-like/interleukin-1 receptor adaptor), and TRIF (the MyD88-independent TLR adaptor that is required for TLR3 and TLR4 signaling pathways) and assessed their contributions to the *CXCL10* promoter activity in the infection +IFNγ conditions (Figure 4D). We did not observe a significant loss in the *CXCL10* promoter activity in the synergistic conditions, when any of these adaptors were silenced compared to that of mock transfected. This indicates that either (i) these pathways are not individually required for increase in IFNγ-mediated *CXCL10* expression during infection and potential redundancy exists between these sensing pathways or (ii) the potential involvement of alternate PRR pathways.

IFNγ has been previously reported to synergize with sterile inflammatory agonists including cytokines such as TNFɑ, IL-1β to induce higher levels of CXCL10 production in various cell types [47,62–65]. This raised the question whether the infection +IFNγ-induced increase in synergistic *CXCL10* expression is in part mediated by secretion of TNF or IL-1β during the course of infection, which in turn synergizes with IFNγ. To test this possibility, HeLa were transfected with siRNA targeting IL-1R or TNFR1 and *CXCL10* promoter activity was measured (Figure 4E). No appreciable decrease in *CXCL10* promoter activity was noticed in either of these cases, if any, silencing of IL-1R significantly increased the response (Figure 4E) indicating that the synergy was not due to an indirect, secondary effect of TNF or IL-1β.

### *C. burnetii* infection specifically augments the expression of multiple IFNγ-induced pro-inflammatory cytokines and is not associated with cell death

To understand if this synergistic effect is specific to CXCL10 or is applicable for other IFNγ-induced immune response genes, we tested the expression of the CXCL10 family members CXCL9 and CXCL11. CXCL9 and 11 share many properties and receptor (CXCR3) as CXCL10 and are regulated by shared and distinct mechanisms [25,66]. Similar to *CXCL10*, we observed a significant increase in expression of *CXCL9, CXCL11* in both HeLa as well as THP1-derived macrophages (Figure 5A-B, 5G-H). In addition, the expression of other prominent early response cytokines including the neutrophil-attracting cytokine IL-8 (chromosome 4) and the pro-inflammatory cytokines IL-6 (chromosome 7) and TNFɑ (chromosome 6) also exhibited a significantly higher expression in *Coxiella-*infected, IFNγ-treated cells (Figure 5C-E), but not other genes (example: *STAT1*) (Figure 5F). These data indicate that at least a subset of IFNγ-induced genes, but not all, across different chromosomal locations exhibit a similar pattern of gene regulation. We analyzed the promoter sequences (∼1000 bp upstream) of these genes (details given in Table 5), for transcription factor binding sites using Find Individual Motif Occurrences (FIMO) in MEME Suite software (https://meme-suite.org/meme/). The details of TF-binding sites and the corresponding nucleic acid sequences with positions relative to transcription start site (+1) are illustrated in Figure 5I. Our annotation is based on (i) motif specifications retrieved from Jasper database (https://jaspar.elixir.no/) (as listed in Table 6), (ii) previously published/annotated/functionally tested by other research groups and (iii) those that conform to consensus sequences (details given in Table 7). In agreement with our luciferase assays indicating the requirement of NF-κB sites in the synergistic expression of *CXCL10* (Figure 3E), the presence of NF-κB binding sequences were observed in the promoter region of genes that exhibited synergistic increase in expression among all those that were tested.

**Figure 5.**
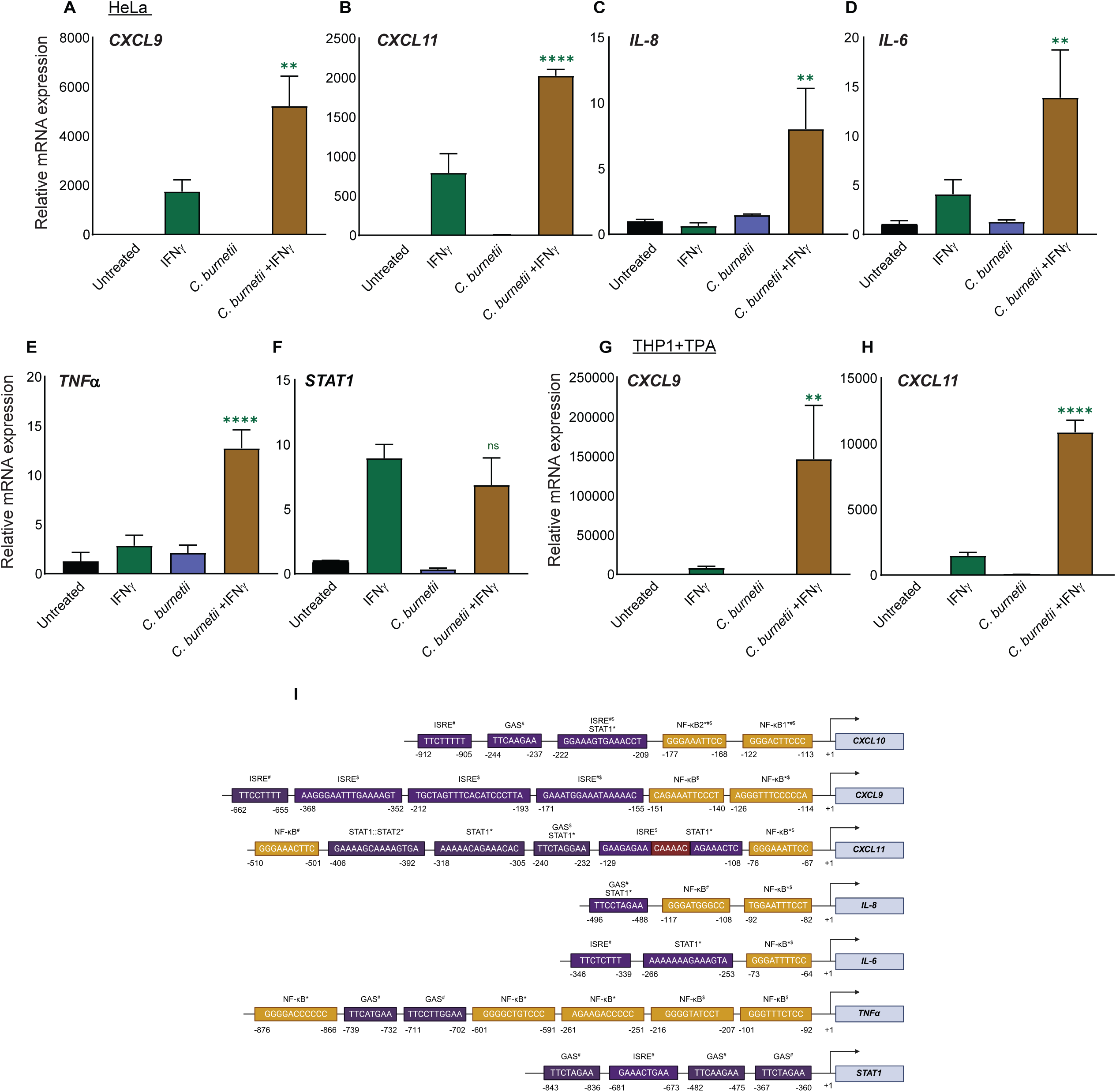
*C. burnetii* infection specifically augments the expression of multiple IFNγ-induced pro-inflammatory cytokines. Relative mRNA expression of *CXCL9* (**A**), *CXCL11* (**B**), *IL-8* (**C**), *IL-6* (**D**), *TNFɑ* (**E**) and *STAT1* (**F**) in HeLa cells infected with *C. burnetii* and supplemented with IFNγ (10 ng/ml) for 24h. Values indicate mean with SD and are representative of three independent experiments (X=3, n=3). Relative mRNA expression of *CXCL9* (**G**) and *CXCL11* (**H**) in the THP1 macrophages in the synergistic condition. The values are normalized relative to GAPDH expression. Statistical analysis was performed using one-way ANOVA with Tukey’s post-hoc test. ** p<0.01, **** p<0.0001 compared to IFNγ alone treatment (green bar). **I**. Annotation of NF-κB, ISRE, GAS and STAT binding regions in the proximal promoter region (−1 to ∼-1000bp) of the above mentioned genes as validated through MEME Suite (https://meme-suite.org/meme/) and published reports of consensus binding sites search or functional annotation. * indicates motifs identified through our JASPAR-MEME-FIMO pipeline for specific transcription factors of interest, ^#^ indicates the sequences that are in close alignment with published consensus binding sites and ^$^ indicates binding sites as annotated and/or functionally mapped by previously published studies. The annotation approaches along with relevant references have been detailed in the materials and methods section. Binding sites for NF-κB, ISRE/GAS/STAT1 and overlap regions are depicted as yellow, blue, and red respectively.

To determine whether the excess cytokine production is a consequence of higher cell death in infected cells treated with IFNγ, we measured cell viability at 3 dpi (the day when cells were supplemented with IFNγ), or 4, or 6 dpi (Figure S3A to C). The cell death observed does not differ significantly between cells treated with IFNγ alone or infection +IFNγ on 3 or 4 dpi suggesting that the high pro-inflammatory cytokine expression and secretion observed (at 4 dpi) is likely not a consequence of differential cell death (Figure S3A-B). By 6 dpi, the viability of HeLa cells is significantly decreased in the IFNγ-treated cells, irrespective of infection (Figure S3C). Further, no disruptive or overt difference in the CCV size or morphology or vacuolar escape/cytosolic release of *Coxiella* was noticed on 5 dpi, between IFNγ-treated or untreated infected cells suggesting loss of CCV integrity or niche (Figure S3D).

### Supplementation of kynurenine does not affect the enhanced promoter activity of *CXCL10* in infected cells treated with IFNγ

The catabolism of tryptophan is initiated by the IFNγ-responsive enzyme IDO1, resulting in the accumulation of kynurenine. Kynurenine subsequently acts as a signaling molecule by binding to the aryl hydrocarbon receptor (AhR), a transcription factor that is involved in modulating various innate and adaptive immune responses [67]. Liu, X et al. reported that, during Mycobacterial infection, Kyn-AhR pathway suppresses CXCL9 and CXCL10 by inhibiting the phosphorylation of STAT1 through suppressor of cytokine signaling 3 (SOCS3) [68]. Transcriptional regulation of *SOCS3* is mediated by the binding of the aryl hydrocarbon receptor (AhR) and its heterodimeric partner, the AhR nuclear translocator (ARNT), to specific xenobiotic response elements (XREs) located within the gene’s promoter region [68–70]. This led to the speculation that kynurenine levels in infection plus IFNγ treatment conditions may be lower than with IFNγ alone, potentially leading to increased CXC chemokine expression due to the loss of a repressive effect. To address this possibility, we treated cells with exogenous kynurenine (100 µM) or the vehicle control DMSO (Figure S4). Supplementation of kynurenine did not significantly affect the high *CXCL10* observed in infection +IFNγ treatment conditions compared to IFNγ alone, suggesting that the synergistic expression of *CXCL10* is not influenced by levels of kynurenine.

### CXCL10 deficiency or sufficiency does not significantly modulate the intracellular replication of *C. burnetii*

Beyond its canonical role as a chemoattractant, CXCL10 has been reported to direct antimicrobial activity against a broad spectrum of pathogens [71–74]. Additionally, CXCL10 interacts with TLR4 beyond its primary CXCR3 receptor to facilitate signaling [75]. Further, the contribution of many ISGs to cell-autonomous immunity against intracellular pathogens, led us to question whether cellular or secreted CXCL10 may contribute to IFNγ-induced restriction of intracellular *C. burnetii* and/or an antimicrobial state in infected and bystander cells through autocrine and paracrine pathways. To test this hypothesis, *CXCL10* expression in HeLa cells was silenced by siRNA prior to IFNγ-treatment and cells were infected with *C. burnetii*-expressing GFP. As evidenced by *Coxiella* fluorescence over several days post-infection (Figure S5A), while knockdown of *STAT1*, the primary ISG-inducing transcription factor of the IFNγ signaling pathway, allowed significantly higher *Coxiella* replication in IFNγ- treated cells compared to that of mock transfection, no significant rescue was noticed in IFNγ-mediated suppression of intracellular *C. burnetii* replication in the case of knockdown of *CXCL10* suggesting that CXCL10 is not required for IFNγ-mediated restriction of *Coxiella*.

Our work had previously demonstrated IDO1 to be a dominant factor for IFNγ-mediated control of *Coxiella* [23,24]. Next to IDO1, CXCL10 is one of the highest transcriptionally induced, STAT1 target genes [76]. IFNγ-induced cell-autonomous immunity involves multiple redundant mechanisms and it is possible that CXCL10 plays a supplementary role in restriction of *Coxiella* in addition to IDO1. To address this, the sufficiency of CXCL10 in influencing intracellular *Coxiella* replication, independent of its IFNγ inducibility, was tested. To this end, we treated HeLa with recombinant human CXCL10 at the indicated concentrations and infected with *C. burnetii* lux after 6 hours (Figure S5B). Intracellular bacterial replication was assessed from 0 dpi to 7 dpi by measuring *Coxiella* luminescence. No significant change was observed in *Coxiella* metabolic activity in the physiological concentration range of 0.1-1 µg/ml of recombinant CXCL10 (Figure S5B). To further assess any possible direct, intracellular as well as signaling effect on intracellular *C. burnetii* replication, we transduced HeLa cells with a Tet-On inducible plasmid that allows expression and secretion of V5-tagged CXCL10 under the control of a tetracycline-inducible promoter (Figure S5C). These cells were treated with increasing doses of Anhydrotetracycline (ATc) and the levels of V5-tagged CXCL10 in the lysates as well as the supernatant was validated by western blotting (Figure S5C). Fold change in the GE of *C. burnetii* at 6 dpi compared to that of 1 dpi indicates that the replication of *C. burnetii* is not significantly affected, and at best, modestly decreased in CXCL10-expressing cells compared to that of control cells (transduced with empty vector), as compared to the significant decrease observed in the IFNγ-treated cells (Figure S5D) suggesting that higher levels of CXCL10 does not prominently influence *C. burnetii* replication.

### IFNγ treatment of *C. burnetii*-infected cells leads to increased detection of extracellular bacteria

Since we did not observe any direct contribution of CXCL10 to cell-autonomous immunity against *C. burnetii* through our approaches using monoculture models, we set out to investigate any potential cell-intrinsic effect of synergistic conditions on intracellular *C. burnetii* replication. IFNγ treatment of infected HeLa cells at 3 dpi reduced intracellular replication of GFP-expressing *C. burnetii* (as measured by *C. burnetii* fluorescence) (Figure 6A). This spurred the question whether synergistic conditions allowed for altered *Coxiella* release or load in the extracellular media. Hence we infected HeLa cells with GFP-expressing *C. burnetii* and exchanged media with fresh media containing IFNγ on 3 dpi and again on 4 dpi (as described in the flow chart in Figure 6B).

**Figure 6.**
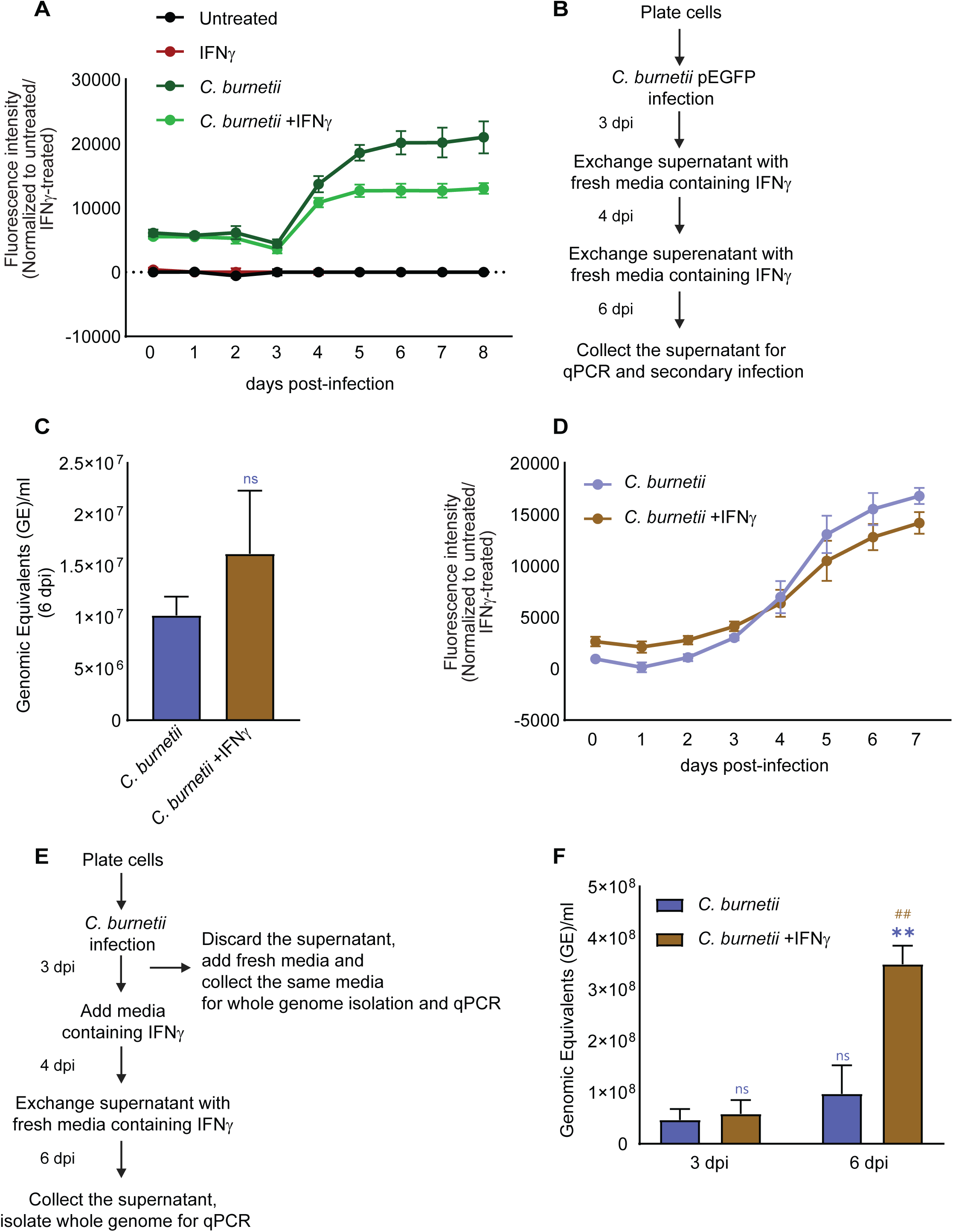
IFNγ-treatment of *C. burnetii*-infected cells leads to increased detection of extracellular bacteria. (**A**) Intracellular replication of GFP-expressing *C. burnetii* in HeLa cells supplemented with IFNγ (10 ng/ml) on 3 dpi for 24 hours. Values indicate mean with SD and are representative of three independent experiments (X=4, n=3). The experimental layout for the measurement of extracellular *C. burnetii* in the synergistic condition and secondary infection is given in panel **B**. The supernatant collected from infected, IFNγ-treated cells on 6 dpi was used for measuring the Genomic Equivalents (GE) of *C. burnetii* (**C**) and for the secondary infection of HeLa (**D**). Values indicate mean with SD and are representative of three independent experiments (X=4, n=3). Statistical analysis was performed using unpaired t-test with equal SD for panel **C**. (**F**) GE measured from whole genomes of *C. burnetii* isolated from the supernatant of HeLa cells infected with *C. burnetii* and supplemented with IFNγ (10 ng/ml) on 3 and 6 dpi (detailed in **E**), as measured through qPCR using *dotA* primers. For panel **F**, values indicate mean with SEM and are the average of three independent experiments (X=4, n=3). Statistical analysis was performed using two-way ANOVA with Tukey’s post-hoc test to compare *C. burnetii* +IFNγ with its corresponding *C. burnetii* alone or between *C. burnetii* +IFNγ of 3 dpi and 6 dpi. ** p<0.01 compared to *C. burnetii*, ^##^p<0.01 compared to *C. burnetii* +IFNγ of 3 dpi.

The supernatant collected on 6 dpi was concentrated, resuspended in media and directly used to measure the Genomic Equivalents (Figure 6C) and parallely, infect naive HeLa cells (secondary infection) to determine the relative levels of viable, infectious bacteria in the supernatants harvested from cells infected with *Coxiella* alone, or infected and supplemented with IFNγ (Figure 6D). The fluorescence-based growth curve indicates that the extracellular bacterial load observed on 6 dpi was viable and infectious and is capable of robust secondary infection (Figure 6D). Extracellular bacterial GE (Figure 6C) as well as *C. burnetii* fluorescence-based growth curves (Figure 6D) indicated near equal or a higher trend of extracellular, infectious bacteria associated with treatment of infected cells with IFNγ (Figure 6C and D). Notably, an increase in the extracellular *C. burnetii* presence on 6 dpi (Figure 6C) coincides with a decrease in the intracellular replication of bacteria in IFNγ-treated cells, at later stages of infection (>5 dpi) (Figure 6A). To further validate these results and to compare against 3 dpi (the day of IFNγ addition when media exchanges ensure that any extracellular bacterial load detected is similar across all infected cells), supernatants from 3 and 6 dpi were harvested sequentially, as described in the flow chart in Figure 6D, whole genome isolated using kits (Figure 6E) and GE measured and compared by qPCR (Figure 6E and F). The extracellular bacterial load between untreated and IFNγ-treated conditions was significantly higher in IFNγ-treated cells on 6 dpi (Figure 6F) compared to that on 3 dpi (the start point), indicating the higher bacterial detection in the synergistic conditions may specifically be attributed to the effects of IFNγ signaling and IFNγ-induced factors. Altogether, these results imply that the extracellular release/load of bacteria is not compromised and in fact, tends to be higher in the synergistic conditions.

### Multiple intracellular pathogens synergize with IFNγ signaling to enhance *CXCL10* expression

To identify if the infection and IFNγ-induced synergistic increase in cytokines is specific for *C. burnetii*, we examined the relevance of this phenomenon to other intracellular pathogens known for their immune evasive strategies. Similar to *Coxiella*, *Salmonella* significantly enhanced IFNγ-induced *CXCL10* expression both at the mRNA and protein level while this augmented expression was abrogated when *Salmonella enterica* serovar Typhimurium (SL1344) was heat-killed (Figure 7A and 7B). Strikingly, infection by the bacterial pathogen *Mycobacterium tuberculosis* (H37Ra) and the parasitic protozoan *Toxoplasma gondii* (Rh decree strain) also led to significantly higher *CXCL10* transcript levels in IFNγ-treated cells (Figure 7C and 7D). This indicates that infection by multiple intracellular pathogens enhance IFNγ-induced CXCL10 production, possibly through a common conserved mechanism.

**Figure 7.**
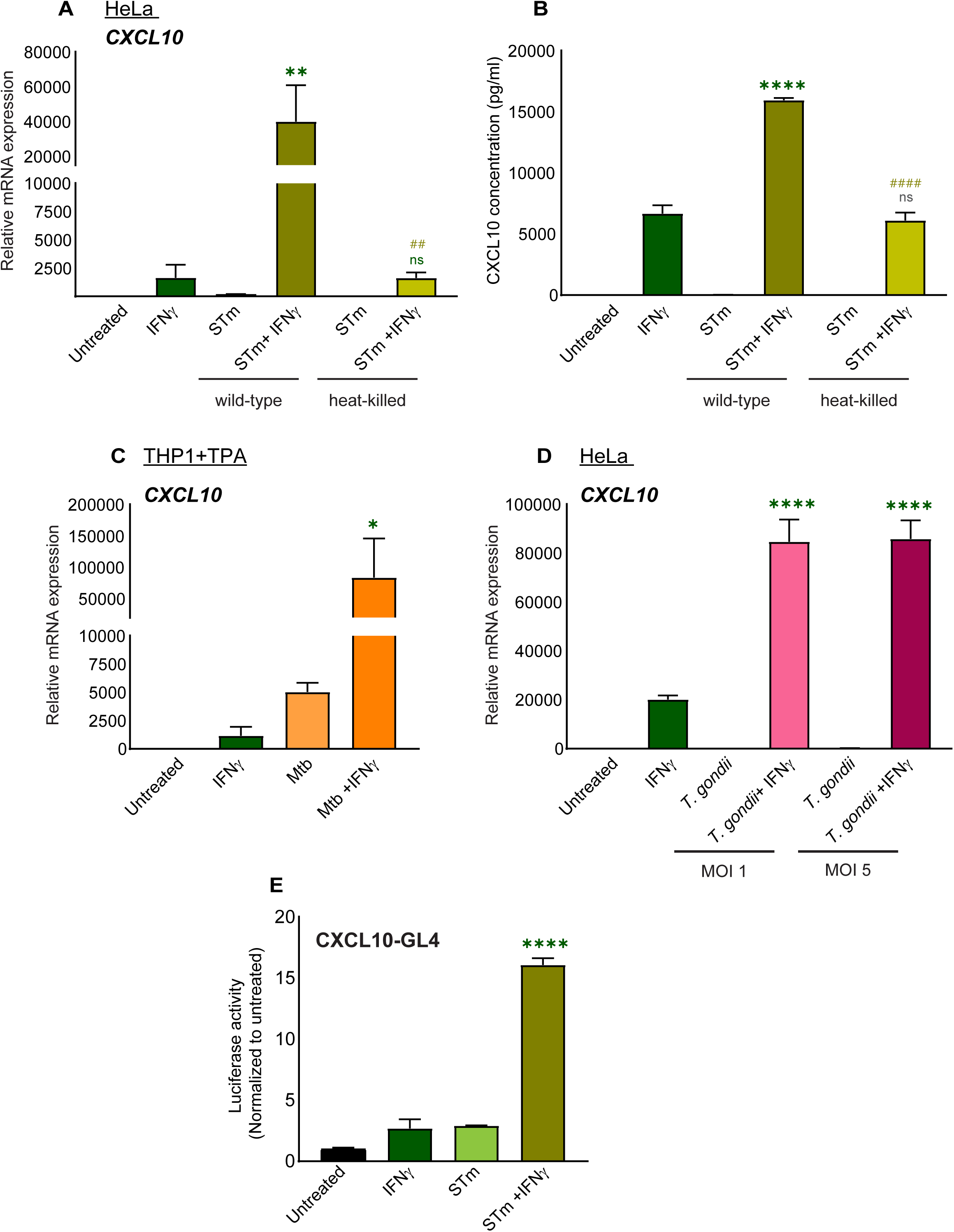
Many intracellular pathogens synergize with IFNγ signaling to enhance *CXCL10* expression. Relative mRNA expression (**A**) and secretion (**B**) of CXCL10 in HeLa cells infected with wild-type or heat-killed *Salmonella enterica* serovar Typhimurium (MOI 20) and treated with 10 ng/ml of IFNγ for 24 hours. Relative mRNA expression of *CXCL10* in TPA-treated THP1 macrophages infected with *Mycobacterium tuberculosis* H37Ra strain (MOI 1) and treated with 10 ng/ml of IFNγ for 6 hours (**C**) and in HeLa cells infected with *Toxoplasma gondii* (at indicated MOIs) and treated with 10 ng/ml of IFNγ for 24 hours (**D**) and values indicate mean with SD and are the representative of three independent experiments (X=3, n=3). The values are normalized relative to GAPDH expression (**A, C** and **D**). Promoter activity of *CXCL10* in HeLa cells transfected with full-length CXCL10 promoter construct (CXCL10-GL4), infected with STm at MOI 20 and treated with IFNγ for 24 hours (**E**). Values indicate mean with SD and are the representative of two independent experiments (X=4, n=2). Statistical analysis was performed using one-way ANOVA with Tukey’s post-hoc test. * p<0.05, **p<0.01, **** p<0.0001 compared to IFNγ, ^##^ p<0.01, ^####^ p<0.0001 compared to wild-type STm +IFNγ.

Similar to *Coxiella*, *Salmonella* infection followed by IFNγ treatment also significantly increased *CXCL10* promoter activity (Figure 7E). We then asked whether the synergy was specific to immunostimulation by IFNγ or whether activation by other proinflammatory cytokines such as TNFɑ, could also mediate this effect. Hence, as a proof-of concept, we treated *Salmonella* infected cells with TNFɑ, instead of IFNγ, for 6 hours and measured *CXCL10* promoter or NF-κB-dependent luciferase gene activity (Figure S6A and S6B). In either case, results indicated a significant decrease in luciferase activity in infected +TNFɑ-treated conditions, indicating that the synergistic relationship between IFNγ and *Salmonella*, (and likely other pathogens) was highly specific and that *Salmonella* still retained its ability to downmodulate other innate immune signaling pathways.

### STAT3 inhibitors decrease the enhanced expression of *CXCL10* in *C. burnetii*-infected infected, IFNγ-treated cells

The persistence of the synergistic expression profile across infection by multiple intracellular pathogens, suggests that the underlying mechanism possibly originates at a point of convergence representing a shared mechanism of pathogenesis. In this context, several intracellular pathogens are known to drive STAT3 phosphorylation and activation as one of the mechanisms to promote an M2 state in macrophages, favourable for their persistent infection and survival and prevent early cell death [77–82]. Indeed STAT3 is one of the genes upregulated during *C. burnetii* infection possibly through IL-6, and is required to support the intracellular replication of *C. burnetii [77,83]*. Hence we tested the possible contribution of STAT3 to the synergistic increase in the expression of *CXCL10* by measuring its promoter activity in the presence of either of the STAT3 inhibitors, C-188-9 and Stattic. C188-9 inhibits STAT3 with high affinity by targeting the phosphotyrosine (pY) peptide binding site within STAT3 SH2 domains (Src homology two domains), thereby interfering with its recruitment to upstream tyrosine kinase-containing complexes, phosphorylation, dimerization and thereby activation [84,85]. Stattic directly targets the SH2 domain of STAT3, irrespective of its phosphorylation state and inhibits STAT3 dimerization and nuclear translocation [86]. Importantly, both the inhibitors were used at the lowest recommended concentration 2.5 µM to avoid any off-target effects. We observed that C-188-9 as well as Stattic, significantly decreased the synergistic increase in *CXCL10* promoter activity, compared to the vehicle control, DMSO (Figure 8A). Notably, no pronounced decrease was observed in the promoter activity in cells treated with IFNγ alone indicating that STAT3 is essential for the synergistic increase in CXCL10 in *C. burnetii*- infected cells treated with IFNγ.

**Figure 8.**
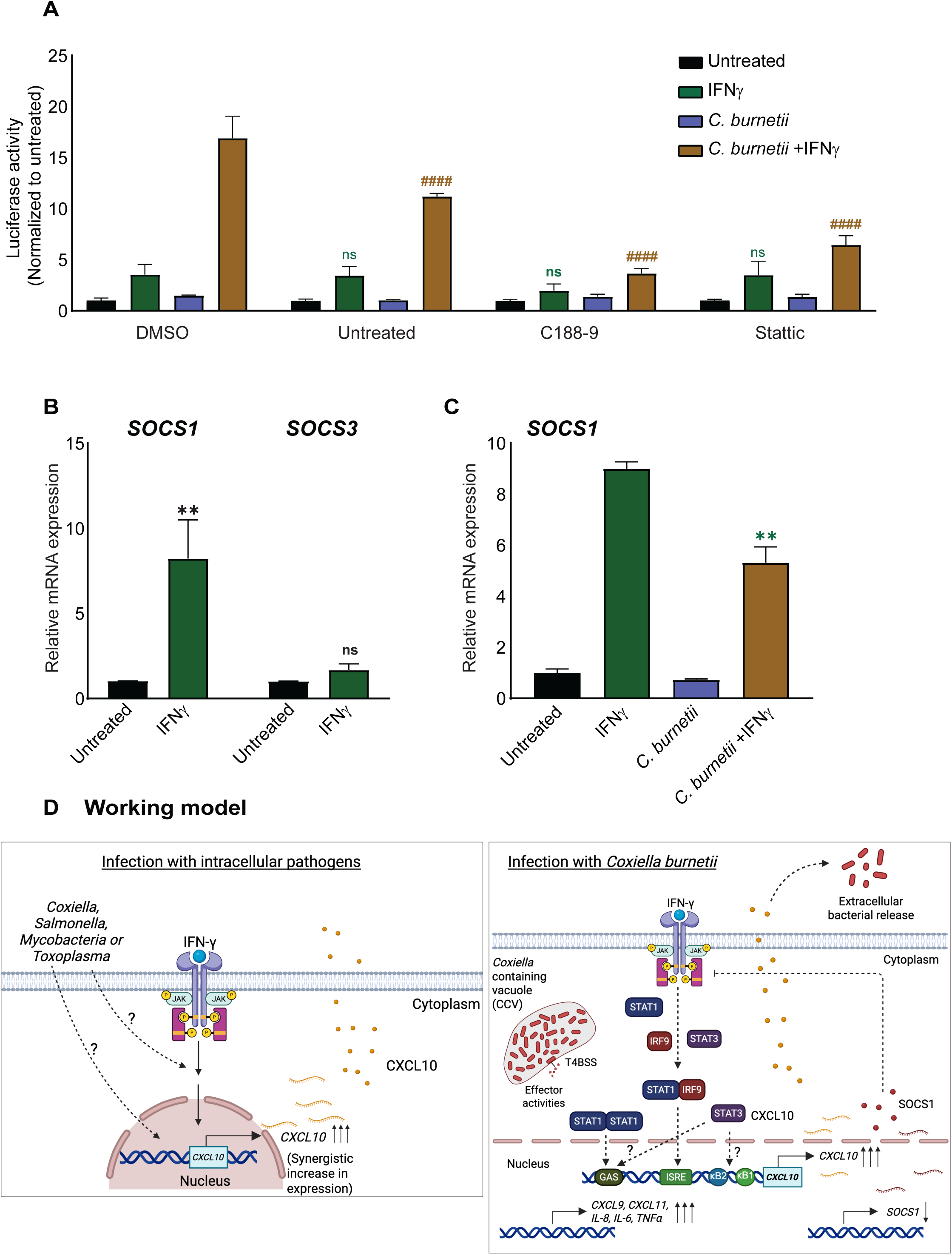
STAT3 inhibitors decrease the enhanced expression of *CXCL10* in *C. burnetii*-infected, IFNγ-treated cells. Luciferase activity of *C. burnetii*-infected HeLa cells transfected with full length CXCL10 promoter construct on 2 dpi, treated with STAT3 inhibitors, C188-9 and Stattic (2.5 µM) on 3 dpi and 6h later, IFNγ was added at 10 ng/ml concentration for 24 hours (**A**). Values indicate mean with SD and are representative of three independent experiments (X=4, n=3). Statistical analysis was performed using two-way ANOVA with Tukey’s post-hoc test. ^####^ p<0.0001 compared to *C. burnetii-*infected cells supplemented with IFNγ (DMSO as vehicle control). (**B**) Relative mRNA expression of *SOCS1* and *SOCS3* in HeLa cells treated with IFNγ at 10 ng/ml for 24 hours. Values indicate mean with SEM and are the average of three independent experiments (X=3, n=3). Statistical analysis was performed using two-way ANOVA with Bonferroni post-hoc test. ** p<0.01 compared to corresponding untreated. (**C**) Relative mRNA expression of *SOCS1* in HeLa cells under synergistic conditions. The values are normalized relative to GAPDH expression, indicate mean with SD and are representative of three independent experiments (X=3, n=3). Statistical analysis was performed using one-way ANOVA with Tukey’s post-hoc test. A working model consolidating the key findings of this study is illustrated in panel **D**.

Pathogens may promote STAT3 signaling not only through direct or indirect activation but also by suppressing negative regulators of STAT activation. Interferon-γ signaling is tightly regulated by multiple negative feedback mechanisms, including the suppressor of cytokine signaling (SOCS) proteins, that attenuate activation of the JAK-STAT pathway and prevent excessive inflammatory responses [87–91]. As the prototypical and potent member of the SOCS family of negative regulators, SOCS1 inhibits JAK-STAT signaling, primarily through direct inhibition of JAK kinase activity while retaining a limited capacity to promote proteasomal degradation of signaling components [87–89,92–94]. SOCS3 also directly inhibits the kinase activity of JAK, but through distinct targeting mechanisms and with an order-of-magnitude reduced potency [93,94]. In addition, SOCS3 has been reported to be a negative regulator of STAT3 as well as CXCL10 as reported in the case of microbial infection [68,95]. To identify the SOCS proteins of relevance in our experimental system, we checked the inducibility of *SOCS1* and *SOCS3* by IFNγ (Figure 8B). Compared to *SOCS3, SOCS1* was found to be significantly induced by IFNγ. Interestingly, synergistic conditions led to a significant decrease in the expression of *SOCS1* gene compared to that in IFNγ treatment alone (Figure 8C) suggesting that higher *CXCL10* in infected +IFNγ treated cells is also correlated with decreased levels of *SOCS1*. Altogether, these results indicate that synergistic expression of *CXCL10* requires STAT3 activity and is associated with reduced *SOCS1* expression.

## Discussion

This study demonstrates that *Coxiella* and a few other intracellular bacterial and parasitic pathogens significantly augment IFNγ-induced expression of a specific subset of genes, including the proinflammatory chemokine, *CXCL10*. Understanding the molecular mechanisms underlying this phenomenon is important to delineate molecular origins of dysregulation in infectious and inflammatory disorders. A working model demonstrating the phenotypes observed, the mechanisms and associated factors identified are illustrated in Figure 8D.

IFNγ has been previously reported to synergize with Influenza virus strain MEM [96], and sterile inflammatory agonists including LPS, cytokines like TNFɑ, IL-1β to induce higher levels of CXCL10 production in various cell types [47,62–65,97]. Multiple overlapping and distinct underlying mechanisms have been reported. In the case of TNFɑ, synergy was observed to be mediated by altered activation and signaling of MAPK transcription factor ERK and reciprocal upregulation of TNFɑRII and IFNGR [63], and increased recruitment of CREB-binding protein (CBP) and RNA polymerase II to the CXCL10 promoter DNA [47]. Synergy was also observed to be regulated by activation of the NF-κB signaling axis in the case of TNFɑ as well as IL-1β in different cell types [62,64,65]; concerted activation of NF-κB and STAT1 by LPS and IFNγ in human neutrophils [97]; NF-κB-dependent and IFNAR-independent mechanisms in the case of influenza virus or dsRNA treatment [96].

Considering this background, a summary of our observations and interpretation of significance of this study, are as follows:

(1) Intracellular pathogens, such as *Coxiella*, *Salmonella, Mycobacteria* and *Toxoplasma* employ highly sophisticated mechanisms to typically inhibit or evade innate and cell-intrinsic immune signaling pathways. However, we observe a synergistic enhancement of at least a subset of ISGs by these pathogens. Importantly, this synergy is not merely additive, which could potentially enable rapid escalation of immune responses at infection sites but also contribute to immunopathology, as seen in chronic inflammatory conditions and cytokine storm syndromes. As witnessed in the case of *Coxiella*, this effect was intrinsically observed in infected, but not by-stander cells, indicating that the amplified cytokine response is not due to soluble, immunostimulatory factor/s.

(2) The augmented expression of *CXCL9, CXCL10, CXCL11, IL-8, IL-6, TNFɑ,* etc., was not observed for few other IFNγ-induced genes (example: *STAT1, SOCS1*) indicating this is a highly regulated phenomenon. Similar to *SOCS1*, it is possible that many other regulators may also be differentially regulated suggesting that the global consequences require further investigation. Interestingly, the genes that exhibit synergism harbor both NF-κB as well as GAS/ISRE binding sites in their promoters suggestive of concerted regulation involving multiple input pathways. - 201G>A, a naturally occurring polymorphism in the promoter of *CXCL10* that enhances its expression is associated with progression of chronic HBV disease in male carriers and more severe Enterovirus-71 (EV71) infection [98,99]. Incidentally, this nucleotide occurs between the ISRE and the NF-κB2 site that were both found to be required for synergistic increase in *CXCL10* expression in the case of *C. burnetii*. Notably, −174G>C and −251T>A polymorphisms in the promoters of the *IL-6* and *IL-8* genes respectively, also enhance gene expression, and here as well, the nucleotides are located between NF-κB and STAT1 binding regions [100–104]. These observations emphasize the importance of the proximal NF-κB and STAT1/ISRE transcriptional elements and their concerted regulation of gene expression.

(3) A functional T4BSS as well as viability were required to observe synergistic increase in CXCL10 levels in IFNγ-stimulated epithelial cells. However, considering the ability of multiple intracellular pathogens to synergize with IFNγ, irrespective of nature of secretion systems, intracellular niche and effector repertoires, we speculate that the synergistic increase in cytokines is not a specific-effector related phenomenon, rather, related to the productive infection process. The divergent mechanism in THP1 cells could, at least in part, arise due to a requirement of “distress/damage” signal or “effector activity-triggered immunity” in the case of epithelial cells which may likely be rendered redundant by the diverse pathogen-sensing mechanisms in THP1 macrophages.

(4) We observed that the synergistic increase in CXCL10 expression required STAT1 and IRF9, but not STAT2. Activation of STAT1 homodimers and canonical ISGF3 complex (STAT1-STAT2-IRF9) are conventionally associated with transcriptional activation of ISGs in Type II and Type I IFN pathways respectively [49,50,52,105,106]. A role of ISGF3 (STAT1, IRF9 and STAT2, likely in unphosphorylated state) has been documented in IFNγ-induced induction of antiviral genes in human cells, similar to a type I IFN response [55,56,107]. Further, the requirement of ISGF3 complexes containing STAT1 and IRF9, but independent of STAT2 has also been described for IFNγ responses [57] which aligns with the observations in our study.

(5) *Coxiella* and other pathogens specifically engage with a pathway that can supplement, but is not sufficient for *CXCL10* induction. Intact NF-κB1 and NF-κB2 sites were specifically required for the augmented *CXCL10* promoter activity in *Coxiella-*infected, IFNγ-treated cells. This is surprising, since *Coxiella* has been demonstrated to inhibit NF-κB activation by distinct effector functions [14,18] and IL-17 signaling-mediated expression of cytokines including TNFɑ, CXCL2, CCL5 [17]. Not just *Coxiella*, infection with *Salmonella* explicitly reduced NF-κB dependent reporter gene expression (Figure S7). This suggests that the NF-κB activating signals necessary for *CXCL10* promoter activation may be distinct, warranting further investigation.

(6) While not examined in this study, it is also possible that synergy is in part, driven by IFNγ- or infection +IFNγ-induced altered chromatin accessibility at target ISG loci driven by epigenetic modifications including histone acetylation and methylation at the promoters, facilitating recruitment of NF-κB and other transcription factors [108–110]. Future studies should also focus on global transcriptomic and epigenetic changes in infected and infected +IFNγ treated cells to glean a comprehensive perspective.

(7) The physiological relevance for why intracellular pathogens may allow for/promote increased production of ISGs is intriguing. One speculation is that increased chemokine expression may recruit a higher number of leukocytes and lead to enhanced dissemination of bacteria from the infected cells to permissive, uninfected cells in the face of an immune stressor such as IFNγ that can also induce cell death. Consistent with this idea, we observed modestly higher bacterial numbers in the supernatants from infected, IFNγ-treated cells, at least in the case of *Coxiella*. Overall, these results suggest that though activation of IFNγ signaling during the course of infection (∼3 dpi) suppresses intracellular bacterial replication, it does not affect exit of viable, infectious bacteria from infected cells.

(8) Although *CXCL10* did not appear to contribute to cell-intrinsic immunity against *Coxiella* in monoculture models using bacterial replication/metabolic activity as a readout, further *in vivo* validation using *CXCL10* knockout models and divergent pathogenic strains is required to delineate its role in host defense and immunopathology.

(9) The data presented in this study indicates that STAT3 and NF-κB are required for infection +IFNγ induced synergistic expression of *CXCL10*. On one hand, STAT3 has been reported to negatively regulate type I IFN signaling and ISGs, including *CXCL9* and *CXCL10*, through sequestration of STAT1 or other possible mechanisms, albeit independent of its role as a transcription factor [111–113]. Indeed, STAT3-deficient tumor cells exhibited enhanced production of ISGs including CXCL9 and CXCL10 [114]. On the other hand, STAT3 is known to bind to GAS sites [115], function in unphosphorylated as well as phosphorylated forms and also form a functional complex with NF-κB [116,117] to positively regulate ISGs expression. STAT1 and STAT3 share high amino acid sequence identity and were analysed to largely bind to similar 9-bp core sequences (optimum binding sites) [118]. Yet, analysis of naturally STAT-regulated genes showed that STAT1 or STAT3 homodimers bind to the promoters of distinct as well as shared transcriptional targets suggesting nuanced specificity [118].

(10) STAT3 inhibitors have emerged as novel therapeutic agents for combating various cancers [119]. While no FDA-approved STAT3 inhibitors are available as yet, small molecule inhibitors are being developed with precise targeting capabilities and some have been employed in clinical phase trials [120]. Given that STAT3 is required for both intracellular pathogen survival, replication [77,83] and IFNγ-mediated exacerbated cytokine production, inhibiting this pathway could potentially offer a dual therapeutic benefit of reducing microbial burden while simultaneously alleviating inflammation-driven pathology.

Altogether, using *C. burnetii* and other intracellular pathogens as model systems, this study provides new insights into the molecular drivers and host-pathogen interactions of infection-induced inflammatory disorders propelling future exploration into the origin of how infection +IFNγ synergy influences systemic infection/pathology and whether it offers a viable window for clinical intervention, and functional cooperation of NF-κB and STAT3 in immune gene regulation.

## Acknowledgements

This work was primarily supported by DBT Wellcome Trust India Alliance Intermediate Fellowship awarded to SG (IA/I/23/2/507001) and intramural funding support from IISER Thiruvananthapuram and Department of Biotechnology- Research Fellowship (Category I) for MVR (DBT/2022-23/IISER-TVM/1881). Our work has been additionally supported by ACORN-AMR fast grant from Ignite Life Science Foundation to SG (ACORN-AMR/2023/003), Start-up research grant (SRG) from the Department of Science and Technology-Science and Engineering Research Board, DST-SERB (now Anusandhan National Research Foundation, ANRF), awarded to SG (SRG/2022/002157), the Department of Biotechnology grant to SG (Indo-UK joint collaborative grant IC-12047(12)/7/2024-BP-IUCA). We are thankful to Dr. Karthik Chandiran for providing training and hands on-help in use of the flow cytometer and Ms. Thanha Aboobacker and Mr. Hrithik Kumar in acquisition and analysis of data. We are grateful for the infrastructural support provided by the Biophysical Instrumentation Facility, Microscopy Core, Flow Cytometry Core and various labs and other Technical and Administrative Staff within the School of Biology (SoB) of IISER Thiruvananthapuram. We express our gratitude to Mr. Tejas Pawar, Ms. Keerthana Baskaran and Ms. Aan Ruth for their valuable support in the establishment of the lab. We also thank Ms. Nayana Harilal, Ms. Tania Elizabeth Manoj, Ms. Persis Prasad, Ms. Rose Mary Joseph, Ms. Vaishnavi Girishan C, Mr. Nandu C K, Ms. Parnika Sahoo, Ms. Ann Mary Alappat, Dr. Kamalakannan Vijayan of IISER Thiruvananthapuram and Dr. Karthik Subramanian (Rajiv Gandhi Centre for Biotechnology (RGCB)) for their valuable time and assistance in the project. We also thank all the members of Host-Pathogen Interaction lab for their valuable help and feedback. We are grateful to Prof. Craig Roy (Yale University), Prof. Jorge Galan (Yale University), Dr. Kamalakannan Vijayan (IISER Thiruvananthapuram) and Dr. Saleem Mohammed (NISER Bhubaneshwar) for providing *Coxiella burnetii* (NMII), *Salmonella enterica* serovar Typhimurium (SL1344), *Toxoplasma gondii* (Rh decree strain), *Mycobacterium tuberculosis* (H37Ra) strains respectively, and Dr. David Proud (University of Calgary) for CXCL10 promoter luciferase constructs. We acknowledge Biorender, Adobe Illustrator and Graphpad Prism platforms for making figures.

## Materials and methods

### Cell lines

THP1 (human monocytic cell line) and HeLa CCL2 cells were used in the study. THP1 monocytes were cultured in RPMI media (Gibco A10491-01) supplemented with 10% heat-inactivated FBS (Corning 35-016-CV). To differentiate into macrophages, THP1 cells were treated with 200 nM of TPA (12-O-Tetradecanoylphorbol-13-acetate, Cell Signaling Technology 4174S). HeLa cells were maintained in DMEM (Gibco 11965126) supplemented with 10% heat-inactivated FBS. For infection studies, DMEM with 5% FBS was used for HeLa cells. All the cells were maintained in the incubator supplied with 5% CO_2_ and maintained at 37°C.

### Coxiella strains

Wild-type Nine Mile Phase II *C. burnetii* strain (RSA439), *icmL*::Tn mutant and luminescence or fluorescence-expressing strains *(C. burnetii* lux*, C. burnetii* pGFP) were used for infection studies (Table 1). All the strains were cultured in 1X acidified citrate cysteine medium 2 (ACCM2), (Sunrise Science 4700-003). For *icmL*::Tn mutant and lux- or GFP-*Coxiella*, Kanamycin (375 µg/ml) was added in ACCM2 as the selection agent. The bacterial strains were incubated at 37°C in a modular chamber supplied with 5% CO_2_, 2.5% O_2_ and 92.5% N_2_. They were cultured for 6 days, centrifuged at 4000 rpm for 15 minutes at 4°C, and the cells were resuspended in DMEM containing 5% FBS. Where indicated, *Coxiella* was heat-killed at 70°C for 30 minutes.

**Table 1.** *C. burnetii* strains used in this study.

| Strain | Antibiotic resistance | Genotype/ Generation | Reference |
| --- | --- | --- | --- |
| Wild-type (wt) Nine Mile phase II (NMII) RSA439 |  |  |  |
| *C. burnetii lux* (generated and gifted by Shawna C. Reed, Roy Laboratory, Yale University) | Kan | Single copy attTN7::KAN-P311-*luxCDABE*-TT genomic insertion via plasmid with *cbu0311* promoter and a terminator cloned into pMiniTn7T-KAN-*luxCDABE* (gift from Paul A. Beare, NIAID, NIH, Montana, USA) | [121] |
| *icmL::Tn C. burnetii* | Kan | Transposon insertion in *icmL.1/dotI (cbu1629)* | [41] |
| *C. burnetii* pEGFP | Kan | *C. burnetii* strain that constitutively expresses GFP | Generated in the Roy lab, Yale University |

### *Coxiella* infection experiments

HeLa cells were plated in 24-well plates at the seeding density of 100,000 cells per well. TPA (200 nM)-treated THP1 cells were plated at the seeding density of 1,000,000 cells per well in a 12-well plate. Next day, the cells were either treated with 10 ng/ml of recombinant human IFNγ (Biolegend, 570208), mouse IFNγ (Biolegend, 575304) or infected with *Coxiella* at the specified MOI and the plates were centrifuged at 500g for 30 minutes prior to incubating the plates in 37°C incubator.

### Synergy experiments

For the synergy experiments, the HeLa cells were plated in a 24-well plate at the seeding density of 50,000 cells per well. The next day, the cells were infected with the *Coxiella* strains as indicated, and the plates were centrifuged at 500g for 30 minutes. Three days post-infection, the media was removed and replenished with fresh media containing IFNγ (10 ng/ml), and the samples were processed for RNA isolation 24 hours later. For the TPA-differentiated THP1 macrophages, the seeding density was 1,000,000 cells per well in 12 well plates. IFNγ (100 ng/ml) was added 2 days post-infection, and samples were collected 6 hours after IFNγ treatment. Where indicated, Chloramphenicol (CAP) at 10 µg/ml was added one day prior to IFNγ treatment.

### Generation of stable cell lines

The coding sequence of human *CXCL10* was cloned into the Tet-on inducible vector pTRIPZ with a linker and a V5 tag through sequence and ligation-independent cloning (SLIC). The details of the primers used for SLIC are listed in Table 2. The cloned vector was transfected in HEK293T cells using Lipofectamine 2000 (Invitrogen, 11668027) at 1.43 µg/well of a 6-plate as per manufacturer’s instructions, and the virus-containing supernatant was used to transduce HeLa cells using polybrene at 8 µg/ml. The cells were allowed to rest for 48 hours and supplemented with fresh media for 24 hours and were subjected to Puromycin selection at the concentration of 1 µg/ml.

**Table 2.**
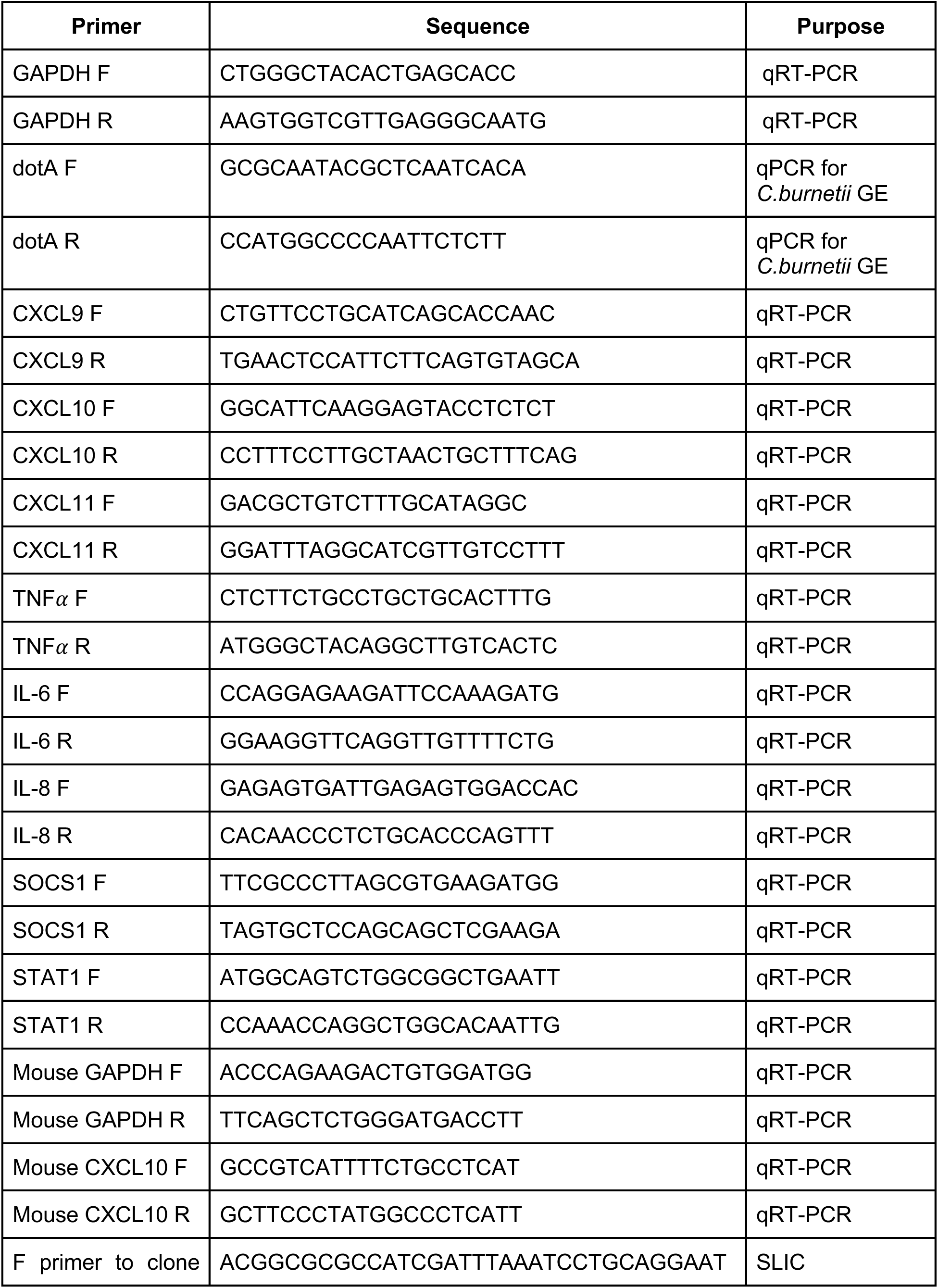
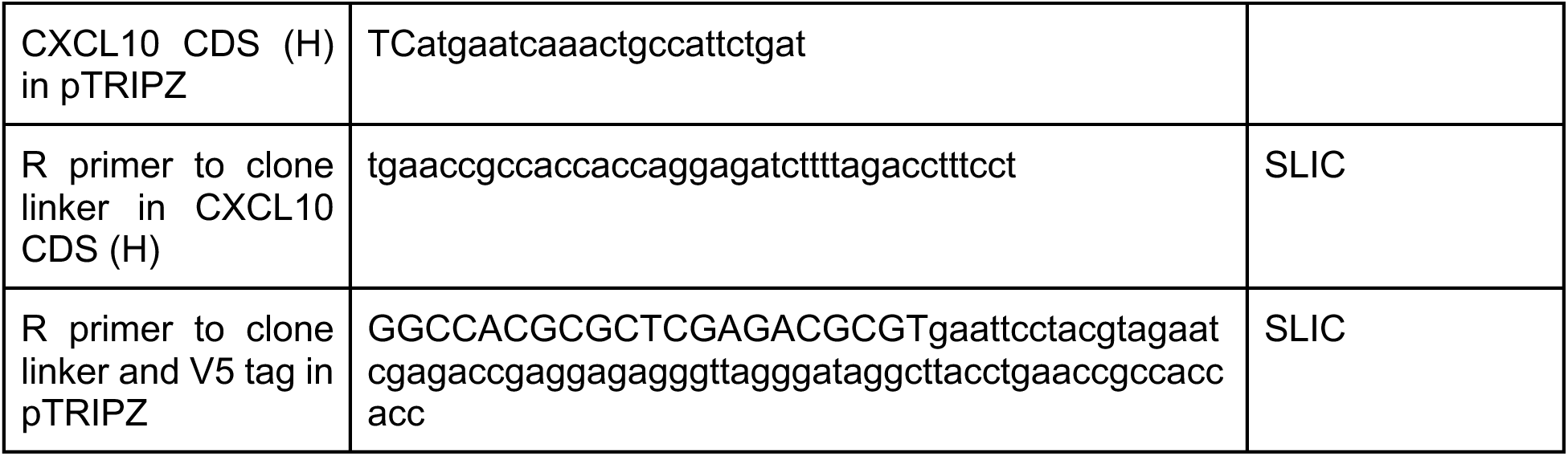
Primers used in this study.

### qRT-PCR

The cells were washed twice with 1X PBS, and RNA was isolated using RNAIso plus (Takara 9109). 1 µg of RNA was converted into cDNA by Takara Primescript RT reagent kit (RR037A), and 37.5 ng of cDNA was used for quantitative real-time PCR. The qPCR reactions were conducted using TB Green® Advantage® qPCR Premix (Takara, 639676) according to the manufacturer’s instructions and the data were collected from Biorad CFX connect real-time PCR detection system. The data were analysed as relative expression, using the ΔΔCt method keeping the uninfected/untreated as calibrator. The list of primers used for qRT-PCR is given in Table 2.

### Dual-luciferase assays

The firefly luciferase reporter pGL4 constructs expressing full length (−875 to +97 bp), truncated (−279 to +97 bp) and mutants of *CXCL10* promoter were generously shared by Dr. David Proud (University of Calgary). HeLa cells were plated at 20,000 cells per well in a 96-well plate, infected with wt *C. burnetii* at MOI 200, transfected with 150 ng of Firefly luciferase plasmid constructs (pGL4) and 50 ng of Renilla luciferase plasmid, pRL-TK on 2 dpi. The cells were treated with IFNγ (10 ng/ml) on 3 dpi and 24 hours later, luciferase activity was measured using Promega Dual-Luciferase assay kit (Promega, E1910), according to the manufacturer’s protocol. For the experiments where STAT3 inhibitor and Kynurenine were involved, infected cells were treated with the inhibitor/Kynurenine at the indicated concentrations on 3 dpi. Six hours post-treatment, IFNγ was supplemented and cell lysates were processed for luciferase assay 24 hours post IFNγ treatment.

**Table 3:** Details of plasmids used in this study.

| Plasmid and Genotype | Notes | Antibiotic resistance | Reference/Source |
| --- | --- | --- | --- |
| CXCL10-GL4 | Firefly luciferase plasmid expressing full length CXCL10 promoter (-875 to +97) | Amp<sup>r</sup> | Gift from Dr. David Proud, University of Calgary [46] |
| tCXCL10-GL4 | Firefly luciferase plasmid expressing truncated CXCL10 promoter (-279 to +97) | Amp<sup>r</sup> |  |
| CXCL10-mCEBPβ1 | Firefly luciferase plasmid expressing CXCL10 promoter with mutated CEBPβ1 binding site (GcGAcAAcGCA) | Amp<sup>r</sup> | |
| CXCL10-mCEBPβ2 | Firefly luciferase plasmid expressing CXCL10 promoter with mutated CEBPβ2 binding site (GtGACTTCaC) | Amp<sup>r</sup> | |
| CXCL10-mκB1 | Firefly luciferase plasmid expressing CXCL10 promoter with mutated NF-κB1 binding site (GtGACTTCaC) | Amp<sup>r</sup> |  |
| CXCL10-mκB2 | Firefly luciferase plasmid expressing CXCL10 promoter with mutated NF-κB2 binding site (GtGAAATTaC) | Amp<sup>r</sup> |  |
| CXCL10-mISRE | Firefly luciferase plasmid expressing CXCL10 promoter with mutated ISRE binding site (GAcAGTGAc) | Amp<sup>r</sup> |  |
| pTRIPZ-EV | Created by annealing the oligos YO-0903 and YO-0904 into the pTRIPZ inducible lentiviral shRNA vector (Dharmacon-Horizon discovery) at Age-I/MluI sites | Amp<sup>r</sup> | Gift from Dr. Brett Lindenbach, Yale University |
| pTRIPZ-CXCL10-V5 | pTRIPZ plasmid expressing CXCL10 gene along with V5 tag, cloned at EcoRI site using SLIC | Amp<sup>r</sup> | This study |

### ELISA

The concentration of CXCL10 on the supernatant was quantified using Peprotech human IP-10 (CXCL10) standard TMB ELISA development kit (Peprotech, 900-T39) according to the manufacturer’s protocol.

### Salmonella infection

Wild-type *Salmonella enterica* serovar Typhimurium (SL1344) was streaked in LB agar from the glycerol stock stored at −80°C and incubated at 37°C overnight. Single colonies were inoculated in plain LB broth (M1245, HiMedia). Overnight culture was further diluted in LB broth at 1:200 ratio. The secondary culture was allowed to reach OD of 0.9 at 600 nm and was used for infection experiments. STm was used to infect the HeLa cells at MOI 20. After 1 hour of infection, the cells were washed twice with 1X PBS and replaced with media with 100 µg/ml Gentamicin. Another hour later, the cells were washed again with 1X PBS twice and maintained in media containing 10 µg/ml Gentamicin. This point is considered as zero hours post-infection. Two hours post-infection, the cells were supplemented with IFNγ at 10 ng/ml, and the cell lysates were processed for RNA isolation after 24 hours of IFNγ addition. For heat-killing, STm was kept in a water bath at 100°C for 30 mins. For luciferase assays, the cells were plated at the seeding density of 20000 cells per well, transfected with the CXCL10-GL4 or NF-κB-RE-GL4 and infected with STm at MOI 20 as mentioned above. After two hours of infection, the cells were treated with IFNγ at 10 ng/ml (for CXCL10-GL4 transfected cells) or TNFɑ at 50 ng/ml (for NF-κB-RE-GL4 transfected cells) and the cell lysates was collected for dual-luciferase assay after 24 hours post IFNγ treatment or 6 hours post TNFɑ treatment.

### Toxoplasma infection

HeLa cells were plated with the seeding density of 100,000 cells per well and infected with *Toxoplasma gondii* (Rh decree strain) at MOI 5 or 10, the next day. After 4 hours of infection, cells were washed twice with 1X PBS and IFNγ (10 ng/ml) was added. Twenty-four hours later, cells were collected and processed for RNA isolation and qRT-PCR for *CXCL10*.

### Mycobacteria infection

TPA-treated THP1 cells were plated with the seeding density of 1,000,000 cells per well and infected with *Mycobacterium tuberculosis* H37Ra strain at MOI 1. The cells were washed with 1X PBS twice after 3 hours of infection. Twenty-four hours later, IFNγ was supplemented at a concentration of 10 ng/ml. Six hours later, cells were collected and processed for RNA isolation and qRT-PCR for *CXCL10*.

### siRNA transfection

HeLa cells were plated with the seeding density of 20,000 cells per well in a 96-well plate and infected with *C. burnetii* at MOI 200 the next day. For the luciferase reporter assays, two days post-infection, cells were transfected with 25 nM of the siRNA using 0.5µl Lipofectamine 2000 per well, followed by IFNγ (10 ng/ml) after 24h. The samples were processed for dual-luciferase assay after 24 hours of IFNγ treatment (4 days post infection). For the *C. burnetii* growth curve experiments, HeLa cells were reverse transfected with 25 nM siRNAs targeting *CXCL10*/*STAT1* as indicated. Forty-eight hours post-infection, cells were infected with GFP-expressing *C. burnetii* (MOI 200) and treated with IFNγ (10 ng/ml), 6 hours post infection. *C. burnetii* fluorescence was measured in the TECAN plate reader (Tecan i-Control 200 pro, Excitation wavelength - 485 nm, Emission wavelength - 538 nm) from 0 dpi till 7 dpi. The details of the siRNA sequences used in this study are given in Table 4.

**Table 4:** Details of SMART pool siRNAs (Dharmacon) used in this study.

| Pool Catalog Number | Gene Symbol | Gene ID |
| --- | --- | --- |
| M-024237-02 | MAVS | 57506 |
| M-024333-00 | TMEM173 | 340061 |
| M-003602-02 | RIPK2 | 8767 |
| M-004769-01 | MYD88 | 4615 |
| M-012833-02 | TRIF | 148022 |
| M-011704-01 | IRF1 | 3659 |
| M-006875-02 | IRF3 | 3661 |
| M-011810-02 | IRF7 | 3665 |
| M-020858-02 | IRF9 | 10379 |
| M-007871-01 | CXCL10 | 3627 |
| M-012064-00 | STAT2 | 6773 |
| M-005188-00 | IL1R1 | 3554 |
| M-005197-00 | TNFR1 | 7132 |
| L-003543-00-005 | STAT1 | 6772 |

### Promoter sequence analysis

The promoter sequence of the genes, 1000 basepairs upstream of the transcription start site, was retrieved from National Centre for Biotechnology Information (NCBI) website (https://www.ncbi.nlm.nih.gov/) (Table 5). Motifs associated with binding of the transcription factors NF-κB, STAT1 or STAT3, as retrieved from Jaspar database (https://jaspar.elixir.no/) are listed in Table 6 and were used for annotation of the promoter sequences of genes of interest. The promoter sequences were scanned for the transcription factor binding site motifs through FIMO in MEME-Suite 5.5.9 software (https://meme-suite.org/meme/tools/fimo). Annotation was based on the following qualifying criteria- (1) q-value<0.05 (q values were obtained from MEME-Suite analyses based on the comparison of each motif with the corresponding promoter sequence, particularly for NF-κB and STAT sites), (2) sequences previously annotated and reported in published literature (example: NF-κB, ISRE and GAS) [65,122–127] and (3) sequences that closely align with the published consensus sites for GAS, ISRE and NF-κB (listed in Table 7).

**Table 5:** Details of the genes and the promoter sequences retrieved from NCBI.

| Gene Name | NCBI Reference Sequence and Gene ID | Chromosome no. | Promoter sequences no. |
| --- | --- | --- | --- |
| *CXCL10* | NC_000004.12 , 3627 | 4 | 76023498 to 76024497 |
| *CXCL9* | NC_000004.12, 4283 | 4 | 76007510 to 76008509 |
| *CXCL11* | NC_000004.12, 6373 | 4 | 76036071 to 76037070 |
| *IL-8* | NC_000004.12, 3576 | 4 | 73739569 to 73740568 |
| *IL-6* | NC_000007.14, 3569 | 7 | 22726200 to 22727199 |
| *TNFα* | NC_000006.12, 7124 | 6 | 31574565 to 31575564 |
| *STAT1* | NC_000002.12, 6772 | 2 | 191014172 to 191015171 |

**Table 6:**
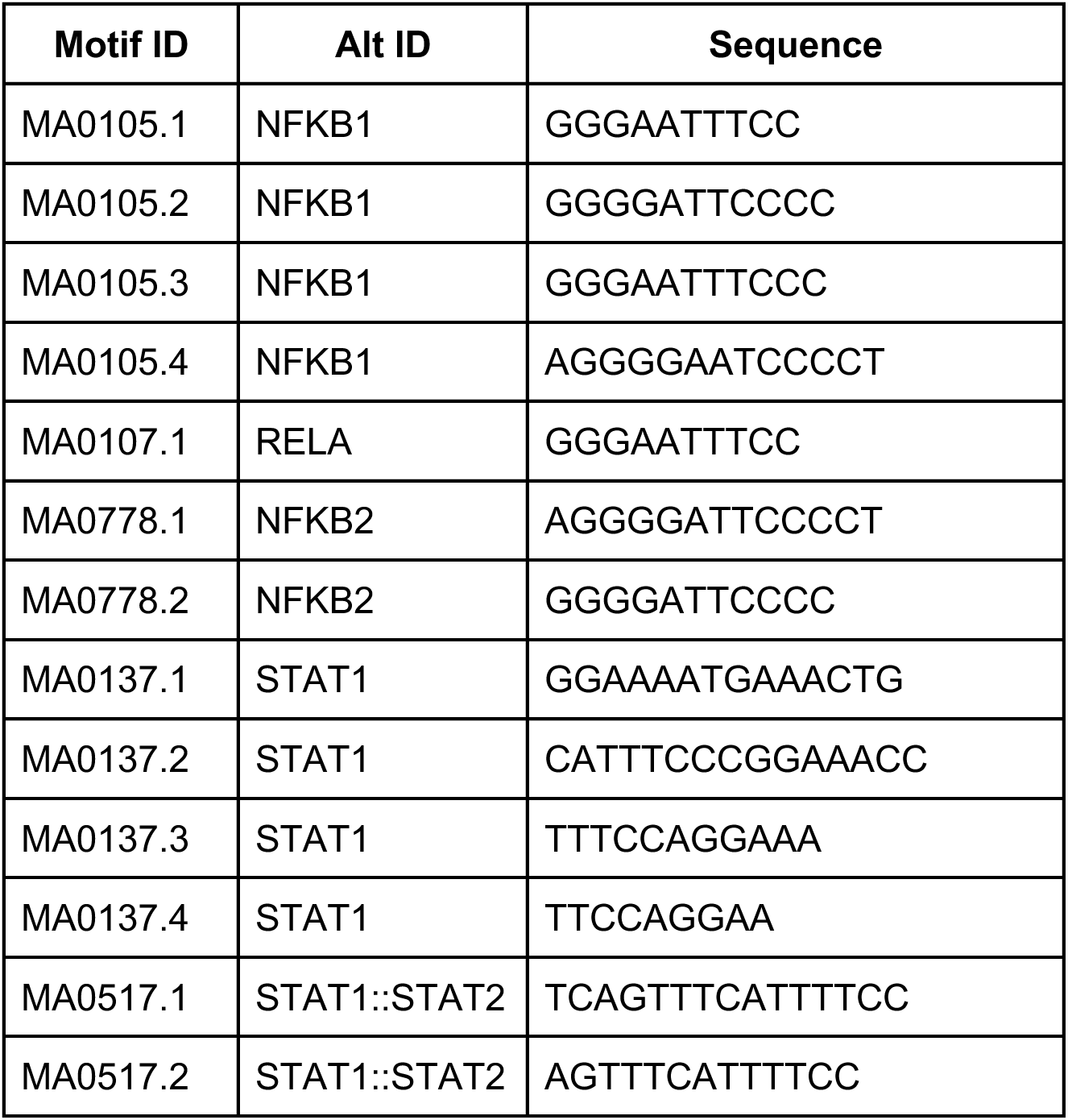
Details of the transcription factor binding sequences retrieved from the Jaspar database, that were used for analyzing the promoter sequence (−1000 to +1) of genes of interest listed in Table 5.

**Table 7:**
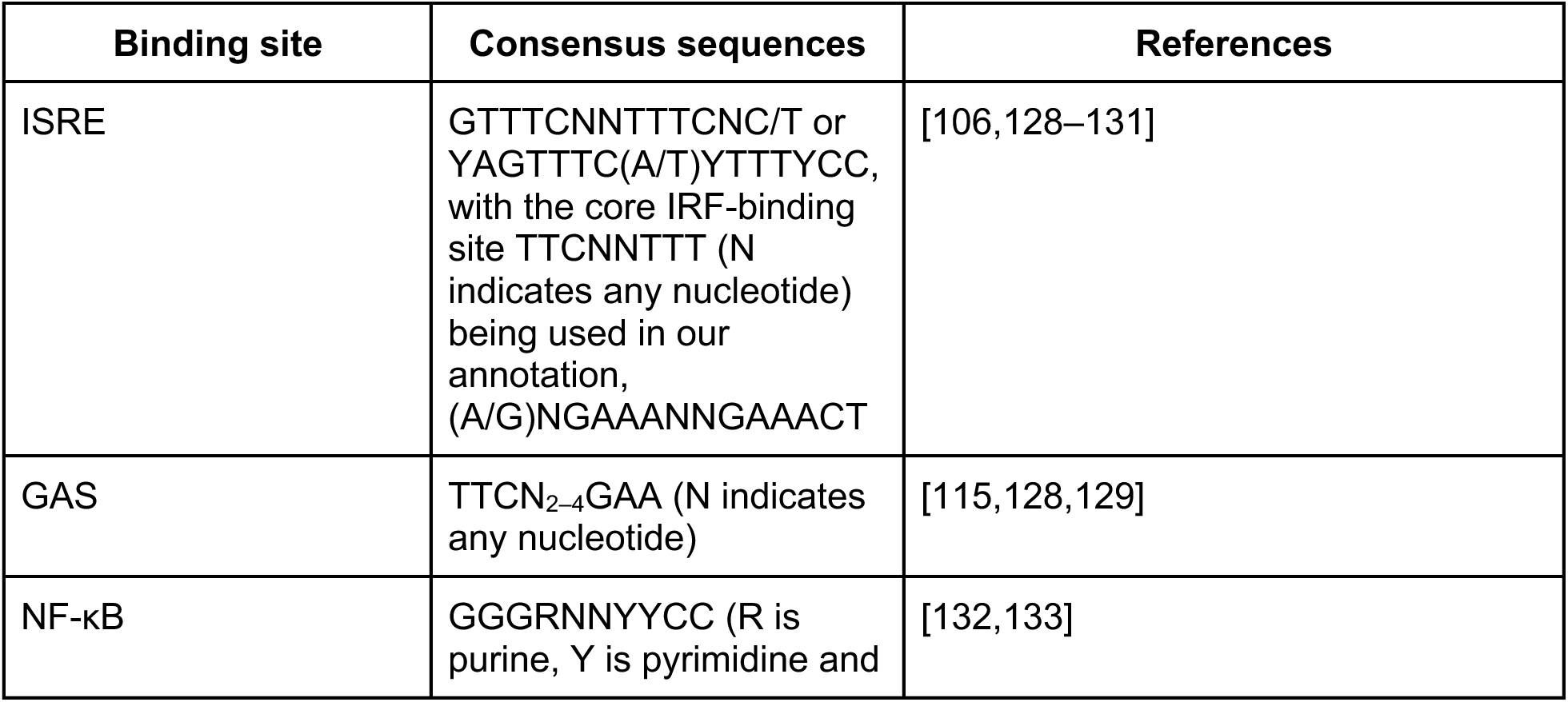
Consensus binding sites of transcription factors of interest.

### Genomic Equivalents measurement of extracellular *C. burnetii*

HeLa cells were plated in a 24 well plate at a seeding density of 50,000 cells per well and infected with *C. burnetii* at MOI 500 the next day. Three days post-infection, cells were replenished with fresh media. The replenished media was collected (3 dpi samples) and the cells were provided with fresh media again with or without IFNγ (10 ng/ml). The supernatant was again exchanged for fresh media with IFNγ on 4 and 6 dpi. The collected supernatants on 3 and 6 dpi were concentrated into a pellet by centrifuging it at 14000 g for 15 minutes, whole genome was isolated using kits (Origin, ODP301-03) and *C. burnetii* genomic equivalents were measured by qPCR using *C. burnetii-*specific *dotA* primers. For experiments involving secondary infection, GFP-expressing *C. burnetii* was used for the primary infection and the pellet from the supernatant collected on 6 dpi was resuspended in 1X PBS and was used to infect HeLa cells plated separately (secondary infection). Intracellular *C. burnetii* replication (secondary infection) was assessed by measuring *C. burnetii* fluorescence using TECAN plate reader (Tecan i-control infinite 200Pro, Excitation wavelength - 485 nm, Emission wavelength - 538 nm). Simultaneously, *C. burnetii* genomic equivalents in the pellet collected from the supernatant of the primary infected cells (6 dpi) was measured by qPCR using *dotA* primers.

### Indirect Immunofluorescence

HeLa cells were plated at a seeding density of 25,000 cells per well in a 24-well plate with cover slips coated with poly L-lysine. The next day, the cells were infected with GFP-expressing *C. burnetii* at MOI 100. Three days post-infection, infected cells were either left untreated or treated with IFNγ at 10 ng/ml. After 24 hours, the cells were fixed with 4% paraformaldehyde (PFA) for 20 minutes followed by washing with 1X PBS and blocking with solution I (0.5% BSA+ 0.2% Saponin+ 1%heat-inactivated FBS in 1X PBS). The coverslips were stained with LAMP1 primary antibody [(DSHB, H4A3), 3 µg/ml] for 1 hour followed by secondary antibody (Goat anti-mouse Alexa Fluor 555, Invitrogen, A28180) and DAPI (4,6-diamidino-2-phenylindole, Sigma, MBD0015) staining (both at 1:2000 dilution) for another hour. The coverslips were mounted using ProLong^TM^ glass antifade reagent (Invitrogen, P36984) on the glass slides and visualized using Zeiss LSM 800 upright confocal multiphoton microscope. The images were processed using Fiji software.

### Intracellular cytokine staining

For intracellular cytokine staining, HeLa cells were infected with either wild-type or GFP-expressing *C. burnetii* for 3 days followed by treatment with or without IFNγ (10 ng/ml) for 24 hours. Eight hours before sample collection, cells were treated with Monensin (0.67 µl per million cells/ml, BD Biosciences D554724). The collected cells were fixed and permeabilized using eBioscience™ Foxp3/Transcription Factor Staining Buffer Set (Invitrogen, 00-5523-00) according to the manufacturer’s protocol, stained with anti-human CXCL10 (Biolegend, 519501) and detected for the intracellular production of CXCL10 through flow cytometry. Data were acquired in BD FACSLyric^TM^ flow cytometer and analyzed using FlowJo software (BD Biosciences).

### Recombinant CXCL10 experiments

HeLa cells were plated at a seeding density of 20,000 cells/well in a 96-well plate and treated with recombinant human CXCL10 (Peprotech, 300-12) or IFNγ (100 ng/ml) the next day. Six hours post-treatment, cells were infected with lux-expressing *C. burnetii* and bacterial luminescence was measured in the TECAN plate reader (infinite 200Pro, Attenuation - none, Integration time - 1000 ms) for a period of 7 days.

### Western blot

HeLa pTRIPZ-*CXCL10*-V5 cells were plated in a 24 well plate at a seeding density of 150,000 cells per well. The cells were treated with Anhydrotetracycline (ATc) at 50 and 500 ng/ml and after 48 hours of ATc treatment, both the cells and supernatant were processed for the detection of CXCL10 through western blot using anti V5-tag antibody (CST, D3H8Q).

### Cell viability assay

The samples meant for positive control were treated with 2.5% Triton-X 100 (HiMedia, MB031). The media was removed and 20 µl of MTT [(3-(4,5-Dimethylthiazol-2-yl)-2,5-Diphenyltetrazolium Bromide), Invitrogen, M6494] solution (5 mg/ml in 1X PBS) was added to the cells and incubated for 4 hours in 37°C. The formazon crystals were solubilized in 100 µl of DMSO and the absorbance at 540 nm was measured using a spectramax plate reader M5.

### Statistical analyses

The data are the representative of at least 3 independent experiments unless mentioned otherwise. The values were analyzed and graphed in Graphpad Prism software version 10. The raw values were analyzed for outliers using ROUT (Robut regression and Outlier removal) method keeping the maximum desired false discovery rate (Q) as 2%. Data that are normally distributed, are analyzed for their significant differences using appropriate one-way or two-way ANOVA followed by the recommended post-hoc test. For the ones that do not follow normal distribution, Kruskal-Wallis test was applied.

### Use of Artificial Intelligence Tools

Generative artificial intelligence (AI) tools, including OpenAI’s ChatGPT and Google’s Gemini, were used during manuscript preparation to assist with language editing, sentence restructuring, refinement of academic writing and relevant academic literature. No AI tools were used to generate research data, perform analyses, interpret findings, or formulate conclusions. All AI-assisted outputs were carefully evaluated and edited to verify accuracy.

## Supplementary figures

**Figure S1.**
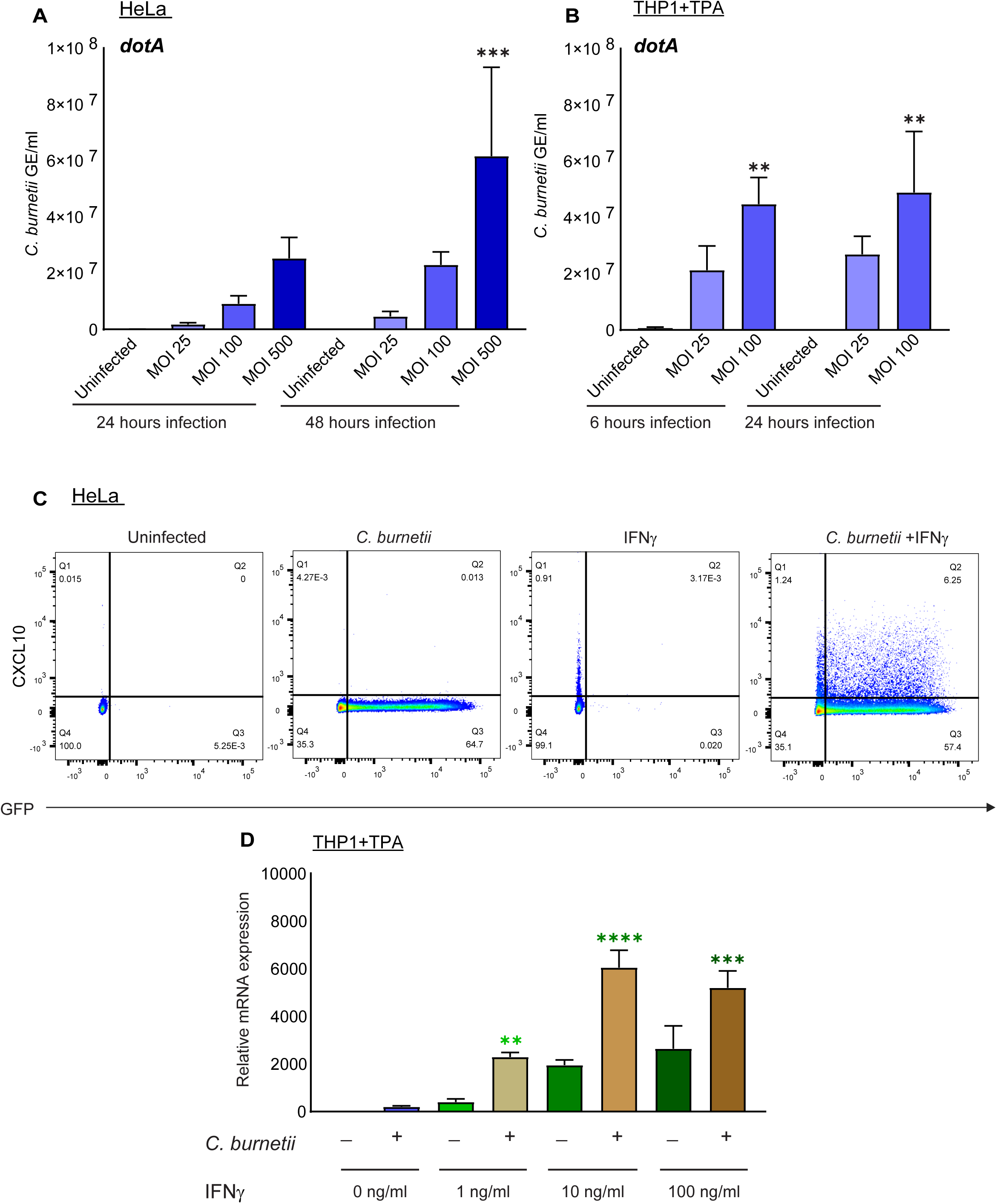
Validation of *C. burnetii* infection, CXCL10 production by infected and by-stander cells and dose-dependent increase in *CXCL10* expression. Genomic equivalents of *C. burnetii* in HeLa (**A**) and TPA-treated THP1 macrophages (**B**) infected with *C. burnetii* NMII at various MOIs for the time points indicated through qPCR using *Coxiella*-specific *dotA* primers. (**C**) HeLa cells were either left uninfected or infected with GFP-expressing *C. burnetii* for 3 days followed by treatment with or without IFNγ (10 ng/ml). Eight hours before sample collection, cells were treated with Monensin (0.67 µl per million cells/ml), stained for CXCL10 and measured by flow cytometer. (**D**) Relative mRNA expression of *CXCL10* in TPA-treated macrophages infected with *C. burnetii* and supplemented with increasing doses of IFNγ (1, 10 and 100 ng/ml), 2 days post-infection and collected after 6 hours. Values indicate mean with SD and are representative of at least two independent experiments. The values are normalized relative to GAPDH expression for panels **A, B** and **D**. Statistical analysis was performed using one-way ANOVA with Tukey’s post-hoc test. ** p<0.01, *** p<0.001 compared to uninfected for **A** and **B** and ** p<0.005, *** p<0.001, **** p<0.0001 compared to its corresponding IFNγ concentration for **D**.

**Figure S2.**
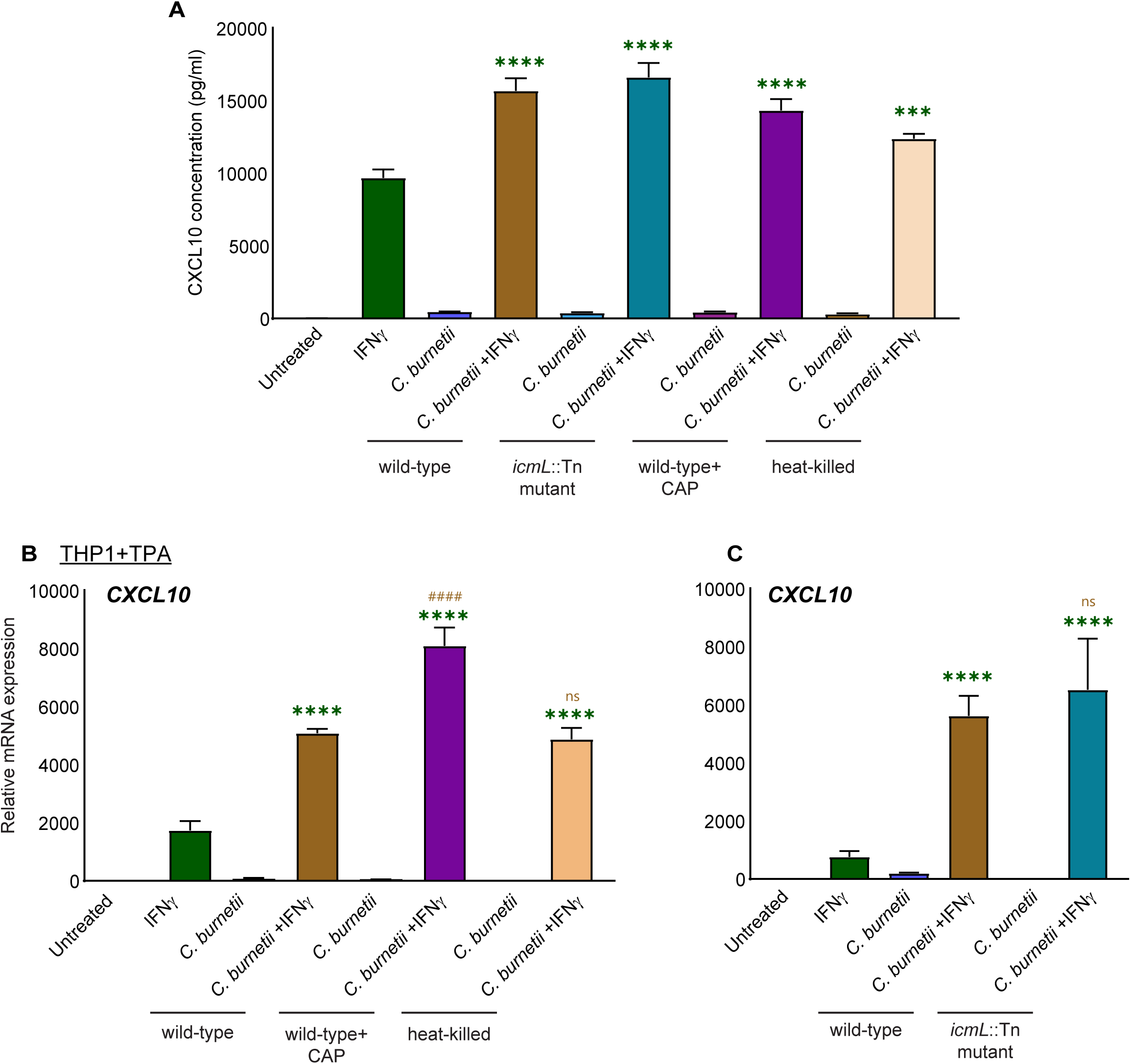
Synergistic increase in *C. burnetii* and IFNγ-induced CXCL10 is not dependent on bacterial viability and T4BSS activity in human macrophages. Protein concentration of CXCL10 in the supernatants of TPA-treated THP1 macrophages in the synergistic conditions. Cells were infected with untreated or heat-killed wild-type or *icmL*::Tn *C. burnetii* strains followed by treatment with chloramphenicol (10 µg/ml) on 1 dpi where indicated. Two days post-infection, cells were treated with IFNγ (100 ng/ml). Samples were collected for ELISA (**A**) and RNA isolation and qRT-PCR (**B** and **C**), 6 hours after treatment with IFNγ. The values are normalized relative to GAPDH expression for **B** and **C**. Values indicate mean with SD and are representative of three independent experiments (X=3, n=3). Statistical analysis was performed using one-way ANOVA with Tukey’s post-hoc test. **** p<0.0001 compared to IFNγ alone (green bar), ^####^ p<0.0001 compared to wild-type *C. burnetii* +IFNγ (brown bar).

**Figure S3.**
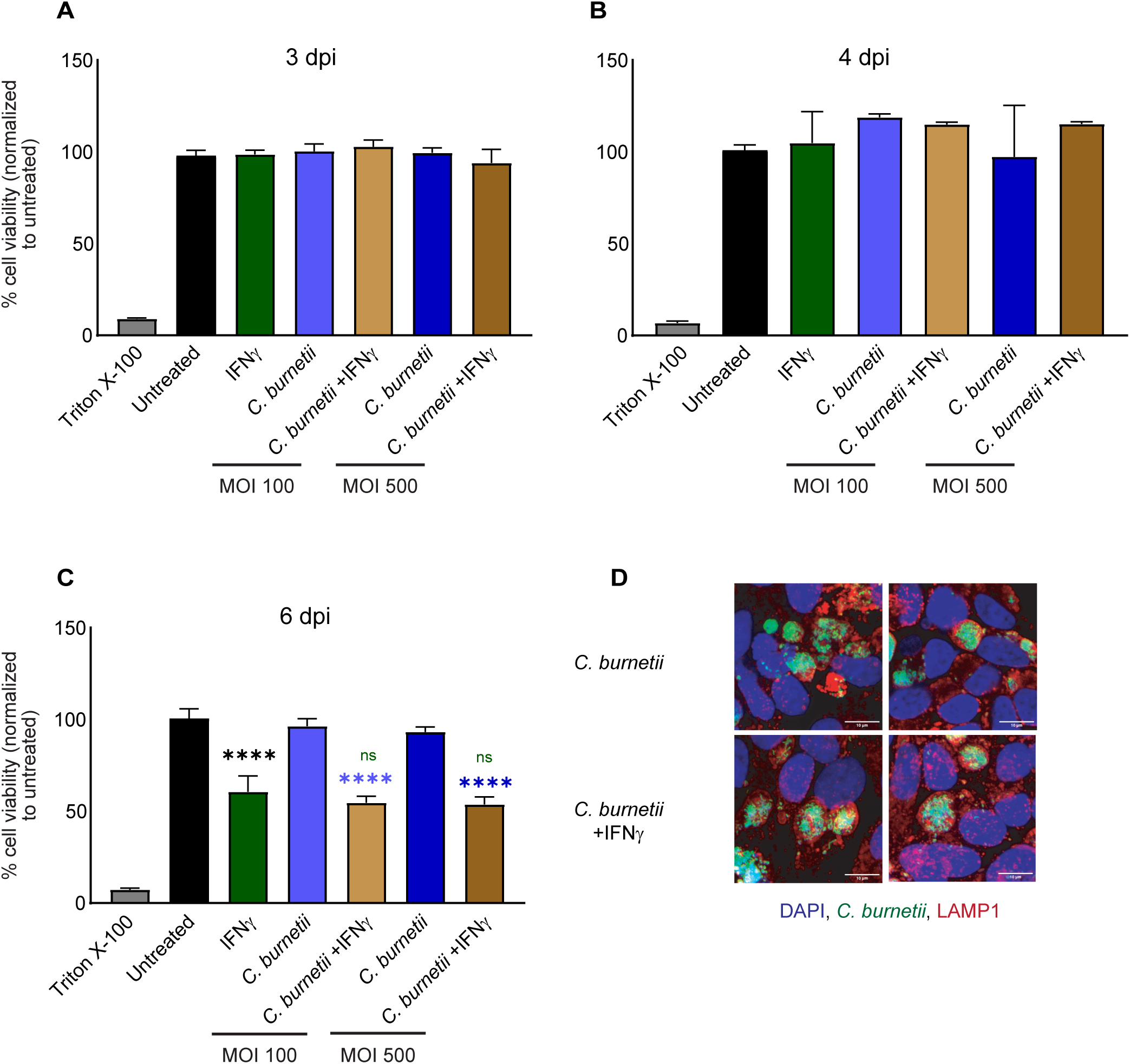
Increase in the expression of pro-inflammatory cytokines in synergistic conditions is not correlated with cell death or gross changes in CCV morphology. Viability of HeLa cells on 3, 4, and 6 days post-infection, either untreated or treated with IFNγ (10 ng/ml) on 3 dpi as measured by MTT assay (**A-C**). Values indicate mean with SD and are representative of three independent experiments (X=4, n=3). Statistical analysis was performed using one-way ANOVA with Tukey’s post-hoc test. ****p<0.0001 compared to the corresponding untreated condition. ***p<0.001 compared to IFNγ-treatment (green bar) or ***p<0.001 compared to *Coxiella* infection alone, of the corresponding MOI (blue bars). (**D**) Immunofluorescent images of HeLa cells on 4 dpi, infected with GFP-expressing *C. burnetii* and treated with IFNγ (10 ng/ml) on 3 dpi, Scale bar: 10 µm. Images are the representative of two independent experiments.

**Figure S4.**
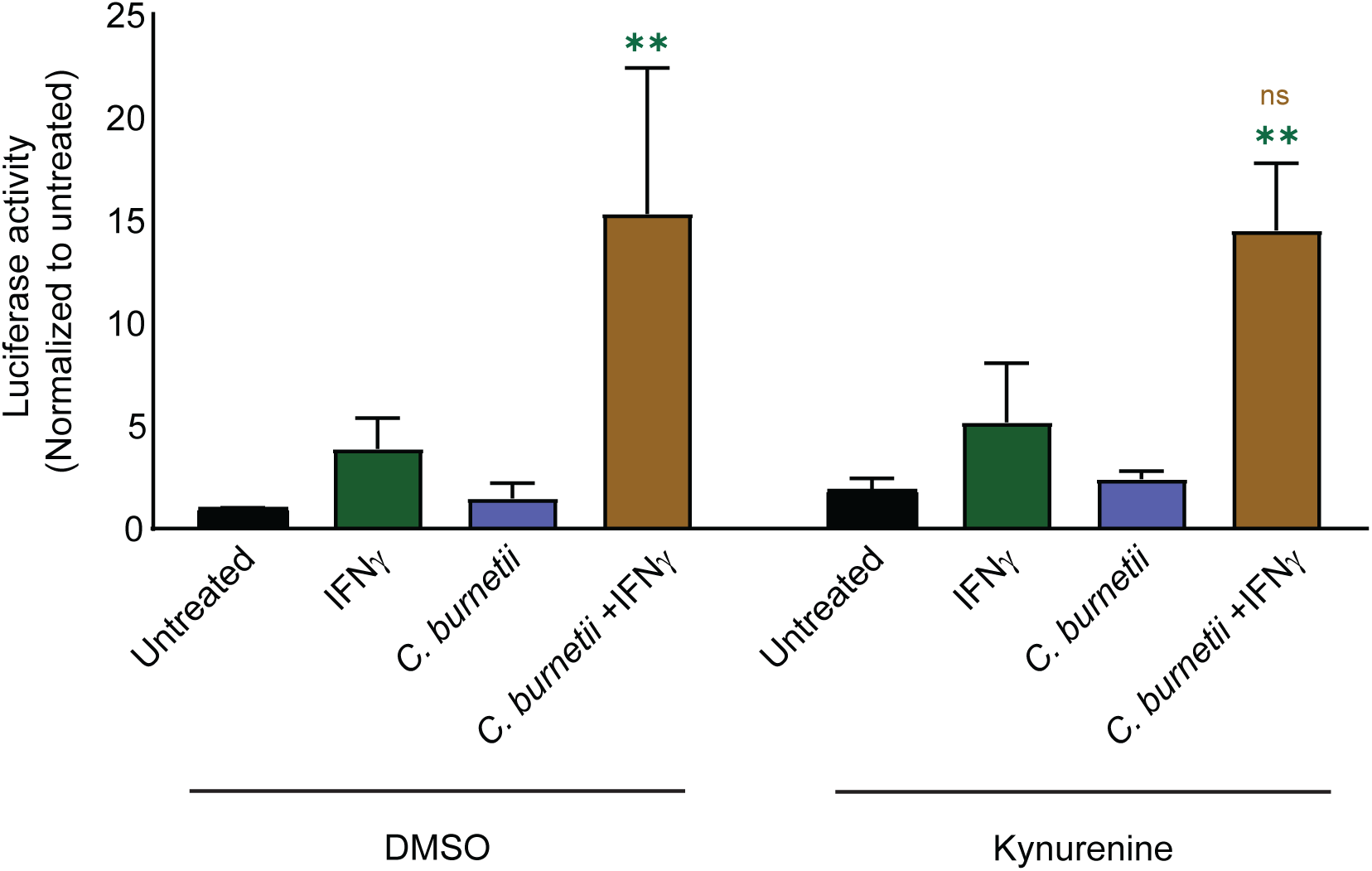
Supplementation of kynurenine does not affect the enhanced promoter activity of *CXCL10* in infected cells treated with IFNγ. Promoter activity of *CXCL10* as measured by dual-luciferase assay in HeLa cells infected with *C. burnetii*, subsequently transfected with full-length *CXCL10* promoter (CXCL10-GL4) on 2dpi, followed by treatment with kynurenine (100 µM) on 3 dpi for 6 hours, followed by supplementation with IFNγ (10 ng/ml) for 24 hours. Values indicate mean with SD and are representative of three independent experiments (X=4, n=3). Statistical analysis was performed to compare luciferase activity of infected +IFNγ treatment with that of the DMSO control (brown bar) or with its corresponding IFNγ control (within the same treatment condition), using one-way ANOVA with Tukey’s post-hoc test. ** p<0.01 compared to IFNγ (green bar) of the corresponding treatment.

**Figure S5.**
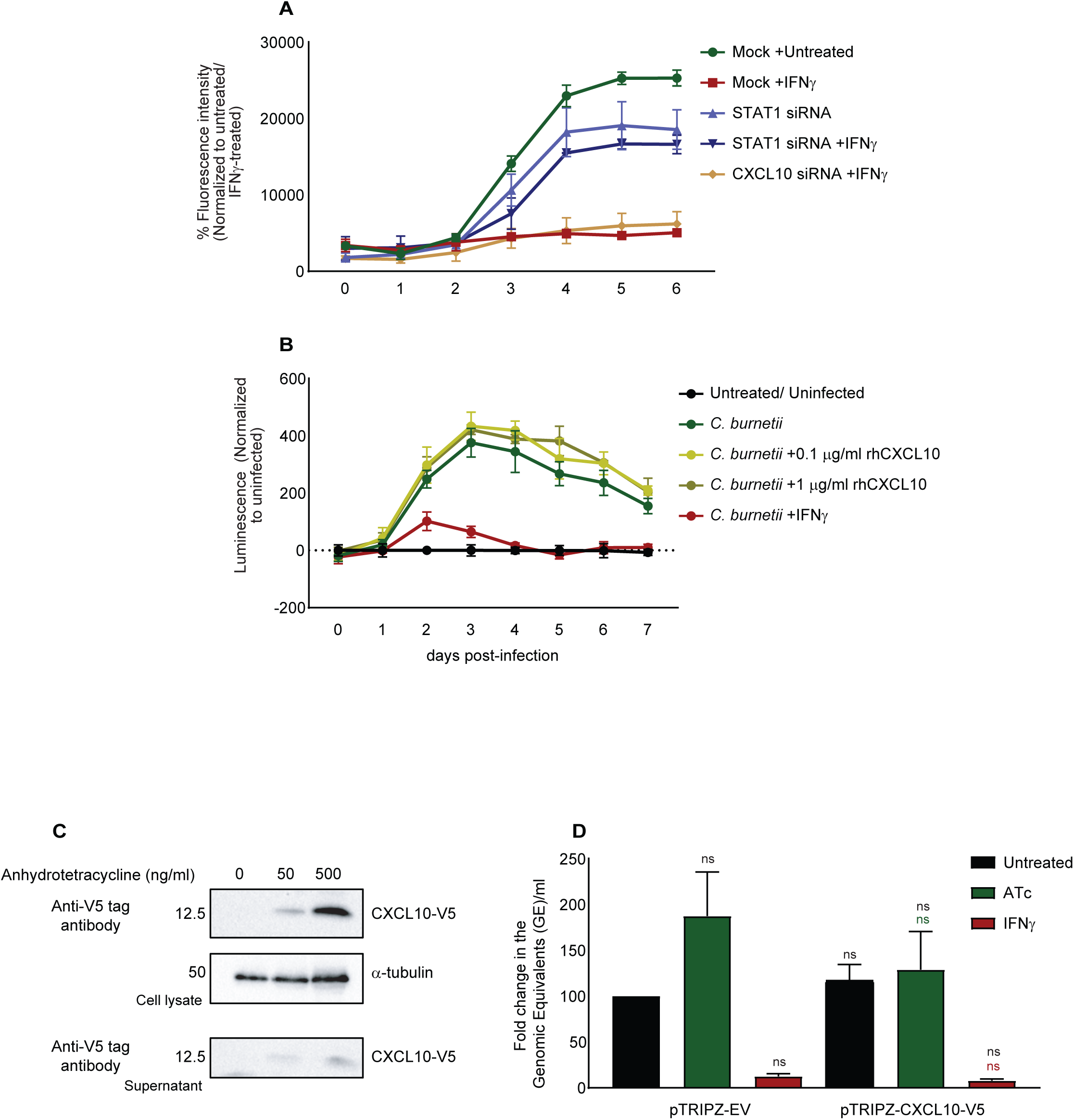
CXCL10 deficiency or sufficiency does not significantly modulate the intracellular replication of *C. burnetii*. (**A**) Intracellular replication of GFP-expressing *C. burnetii* as measured by the percentage fluorescence intensity (normalized to Mock +Untreated) on 5 dpi in HeLa. Cells were reverse transfected either with mock or the indicated siRNAs for 48 hours, infected with GFP-expressing *C. burnetii* and supplemented with IFNγ (10 ng/ml), 6 hours post infection. Values indicate Mean with SD and are the representative of three independent experiments (X=3, n=3). Statistical analysis was performed using one-way ANOVA with Tukey’s post-hoc test. **** p<0.0001 compared to Mock +Untreated, ^####^ p<0.0001 compared to Mock +IFNγ. (**B**) HeLa cells were treated with the indicated concentrations of recombinant human CXCL10 or IFNγ and infected with *C. burnetii* lux, 6 hours post rhCXCL10/IFNγ treatment and the intracellular replication of the bacteria was assessed by measuring the luminescence over a period of 7 days. Values indicate mean with SD and are representative of three independent experiments (X=4, n=3). (**C**) Validation of the expression of CXCL10 in the cell lysate as well as supernatant of HeLa-pTRIPZ-*CXCL10*-V5 cell line upon inducing them with Anhydrotetracycline (ATc) at the indicated concentrations for 48 hours through western blot (representative of two independent experiments). (**D**) Fold increase in the Genomic Equivalents of *C. burnetii* in the ATc/IFNγ treated pTRIPZ-EV and pTRIPZ-*CXCL10*-V5 cells on 6 dpi compared to that of 1 dpi. Values indicate mean with SEM, normalized to pTRIPZ-EV untreated and are the average of 5 independent experiments (X=5, n=3). Statistical analysis was performed using non-parametric one-way ANOVA (Kruskal-Wallis test) with Dunn’s post-hoc test.

**Figure S6.**
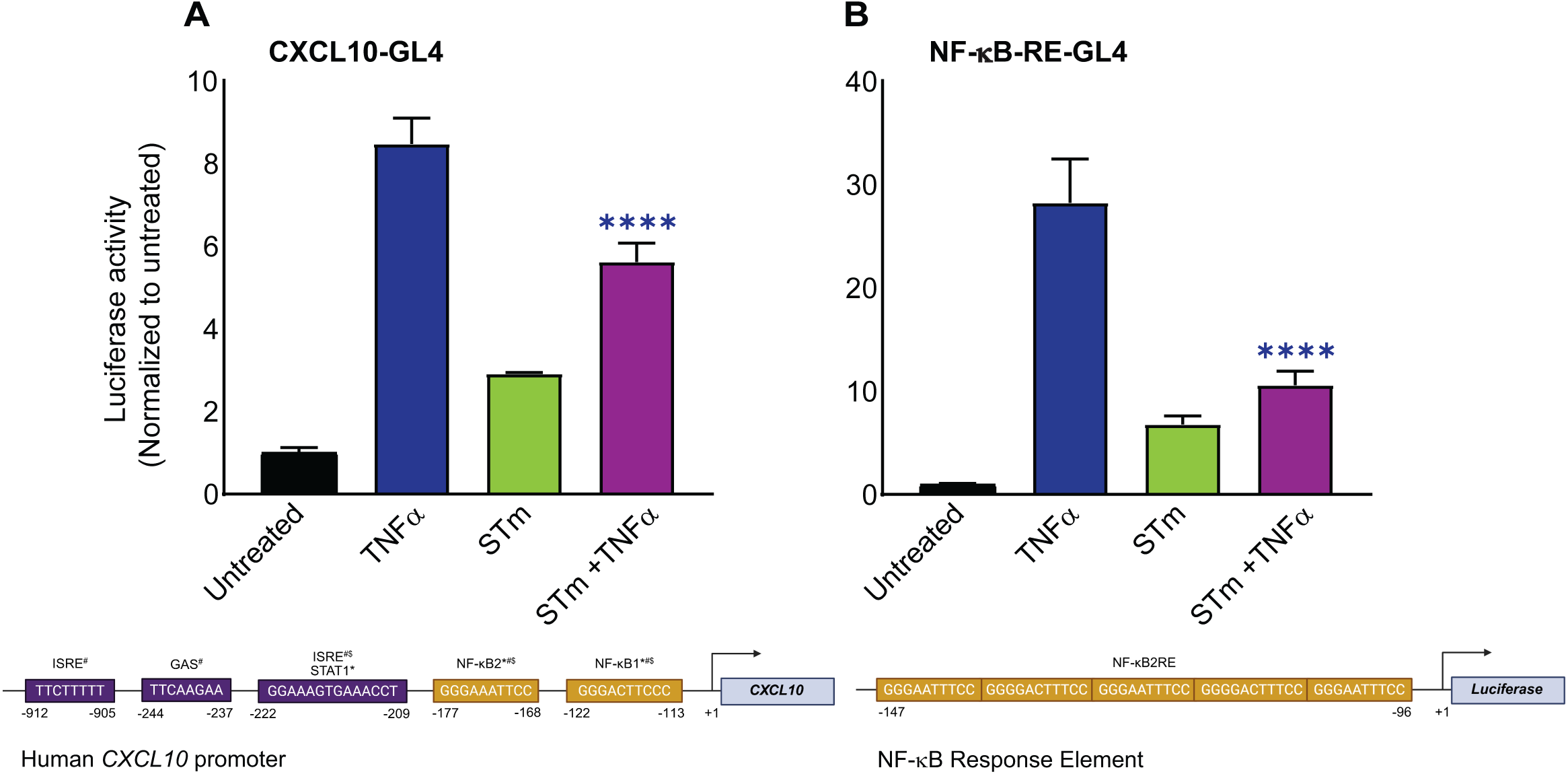
*Salmonella* infection downregulates *CXCL10* promoter- and NF-κB- dependent luciferase activity upon TNFα treatment. *CXCL10* promoter- (**A**) and NF-κB- dependent (**B**) luciferase activity in HeLa cells transfected with full-length CXCL10 (CXCL10-GL4) or NF-κB response element containing promoter construct, infected with STm at MOI 20 and treated with TNFɑ for 6 hours. Values indicate mean with SD and are representative of two independent experiments (X=4, n=2). Statistical analysis was performed using one-way ANOVA with Tukey’s post-hoc test. **** p<0.0001 compared to TNFɑ treatment (blue bar). The details of the prominent promoter elements of the corresponding plasmids are depicted below the corresponding figure panels.

